# The mammalian rod synaptic ribbon is essential for Ca_v_ channel facilitation and ultrafast fusion of the readily releasable pool of vesicles

**DOI:** 10.1101/2020.10.12.336503

**Authors:** Chad Paul Grabner, Tobias Moser

## Abstract

Rod photoreceptors (PRs) use ribbon synapses to transmit visual information. To signal ‘no light detected’ they release glutamate continually to activate post-synaptic receptors, and when light is detected glutamate release pauses. How a rod’s individual ribbon enables this process was studied here by recording evoked changes in whole-cell membrane capacitance from wild type and ribbonless (RIBEYE-ko) rods. Wild type rods created a readily releasable pool (RRP) of 92 synaptic vesicles (SVs) that emptied as a single kinetic phase with a τ < 0.4 msec. Lowering intracellular Ca^2+^-buffering accelerated Ca_v_ channel opening and facilitated release kinetics, but RRP size was unaltered. In contrast, ribbonless rods created an RRP of 24 SVs, and lacked Ca_v_ channel facilitation; however, Ca^2+^ channel-release coupling remained tight. The release deficits caused a sharp attenuation of rod-driven light responses measured from RIBEYE-ko mice. We conclude that the synaptic ribbon facilitates Ca^2+^-influx and establishes a large RRP of SVs.

## Introduction

Animals use their sensory systems to interact with and navigate through their environment, and sensory maps are created for this purpose. This is especially true for vision, where perception of a real-world scene invariably asserts the location of objects in space. The building blocks for visual percepts originate from a visual field that is often in motion, and can vary greatly in light levels. Therefore, vertebrates have evolved complex processes that stabilize the eyes on the visual field (Straka and Baker, 2013), focus images on the back of the eye, and transform light of varying intensities into neural signals (Rivlin-Etzion et al., 2018).

The neural retina lines the concaved inner surface of the eye, and it is a thin, multi-layered network. The outer most layer is a dense lawn of photoreceptors (PR >1 × 10 mm) (Grunert and Martin, 2020). Each PR has a photosensitive outer segment, and at its opposite pole a single synaptic terminal forms in the first synaptic layer of the retina, the Outer Plexiform Layer (OPL). There are two classes of PRs in the outer retina: rods and cones. They make very distinct connections to the inner retina (Behrens et al., 2016; Tsukamoto and Omi, 2016, 2017), the rods are ~10^3^-fold more sensitive to light than cones (Cao et al., 2014; Grimes et al., 2018), and in most mammals rods outnumber cones by 30:1 (Grunert and Martin, 2020). In the primate retina, the rod-dominant region is referred to as ‘peripheral retina’, occupies 95 % of the retinal area, and supports low acuity vision (e.g., peripheral vision) (Sterling, 2013); whereas, the cone-dominant ‘central retina’ supports high acuity, color-sensitive vision (Thoreson and Dacey, 2019).

Both rods and cones form ‘ribbon synapses’ that are named after the electron-dense plate that projects from the presynaptic AZ into the cytoplasm. A subset of vertebrate sensory neurons express the protein RIBEYE, which is localized to ribbons (reviews, Lagnado and Schmitz, 2015; Moser et al., 2019). Deletion of RIBEYE eliminates synaptic ribbons (Maxeiner et al., 2016). So far, studies carried out on ribbonless bipolar cells and auditory inner hair cells have not identified a unifying role for the ribbon. Evoked release from ribbonless bipolars was greatly reduced, though Ca^2+^ currents were normal (Maxeiner et al., 2016). In contrast, ribbonless hair cells showed a milder impairment in exocytosis (Becker et al., 2018; Jean et al., 2018), and they produced well defined substitute AZs that may be better at compensating for loss of ribbons (Jean et al., 2018). These studies suggest that only the synaptic ribbons in the inner retina have a prominent role in vesicle fusion, where ribbons are proposed to physically shorten the coupling of Ca_v_ channels to SVs (Maxeiner et al., 2016). How the ribbon influences transmitter release from mammalian PRs has not been tested directly, and in addition, relatively little is known about the biophysics of exocytosis from mammalian PRs.

Mammalian rods express a single, large horseshoe-shaped ribbon that surrounds one or two ON-bipolar dendrites that are on average ~250 nm away (for mouse; Hagiwara et al., 2018). In the dark rods are maximally depolarized to produce a steady influx of Ca^2+^ that drives the continual turnover of SVs. This keeps synaptic glutamate high enough to activate the postsynaptic inhibitory mGluR6→TRPM1 pathway in the rod ON-bipolar dendrite (Koike et al., 2010), which equates to the ‘dark signal’. A weak flux of photons is sufficient to hyperpolarize the rod and momentarily slow exocytosis to create the ‘light signal’ (for review, Field and Sampath, 2017). Mathematical models have predicted that a rod ribbon needs to achieve a release rate ≥ 40 SVs-s^−1^ in the dark to keep the rod ON-bipolar pathway continually activated (Rao-Mirotznik et al., 1998; Hasegawa et al., 2006).

In the current study high resolution measurements of evoked SV turnover were made directly from mouse rods. The results demonstrate that the rod ribbon creates multiple, uniformly primed SV release sites. Surprisingly, the Ca_v_1.4 channels activated rapidly (~200 μs) and exhibited unique forms of facilitation that influenced ultrafast release rates. In addition, the channels supported continual Ca^2+^ entry which drove the steady turnover of 288 SVs-s^−1^. These features were dependent on the ribbon, as ribbonless rods formed a much smaller RRP, and lacked Ca_v_ channel facilitation. The study provides experimental results that support longstanding proposals on the function of the rod ribbon synapse, and for the surprisingly rapid release kinetics, we discuss how this can contribute to rod dependent vision.

## Results

### Super-resolution readout of SV turnover at an individual rod ribbon

The majority of rod somata reside in the outer nuclear layer (ONL) and send a spindly axon to their singular presynaptic terminal, which contains an individual synaptic ribbon in the OPL. The minority of rod somata that are not positioned in the ONL, but within the OPL, lack an axon and contain the synaptic ribbon within the soma compartment (see illustration in Figure 1A). In the current study, the axonless rods were targeted to optimize voltage control of the ribbon, and more rapidly activate Ca_v_ channels (Hagiwara et al., 2018). A profile of evoked Ca^2+^-current (I_Ca_) measured from a rod filled with an intracellular concentration of 10 mM EGTA is presented to highlight the resolution the signals (Fig. 1C). To measure the fusion of SVs in response to stimulated Ca^2+^-entry, the addition of SV membrane to whole-cell membrane capacitance (C_m_) was monitored with the sine wave based, lock-in amplifier method (Lindau and Neher, 1988). The example in Figure 1D illustrates a rod’s response to a 9 msec depolarization (I_Ca_ presented in Fig. 1C), which shows an ~ 4 fF C_m_ jump (ΔC_m_) that was 13-fold above the standard deviation in baseline C_m_ (~300 aF). The other lock-in amplifier outputs: membrane conductance (G_m_) and series conductance (G_s_) were not affected. The ΔC_m_ was equated to the number of fusing SVs with a conversion factor of 35.4 attoFarads (aF = 10^−18^F) for a 32.5 nm diameter SV (see Methods); thus, a ΔC_m_ of ~ 4 fF is equivalent to ~ 110 SVs. Inspection of the lock-in signals, on an absolute scale, to a series of stimulation epochs further emphasizes the specificity of the ΔC_m_ that followed each stimulation (Fig. 1E). The quantification of ΔC_m_ over baseline and stimulation segments is illustrated in Figure 1H, and the results are plotted over the time course of the experiment (Fig. 1I). The summary of evoked ΔC_m_ plotted as a function of stimulation duration reveals how rapid the rod fused > 100 SVs (Fig. 1J).

**Figure 1.**
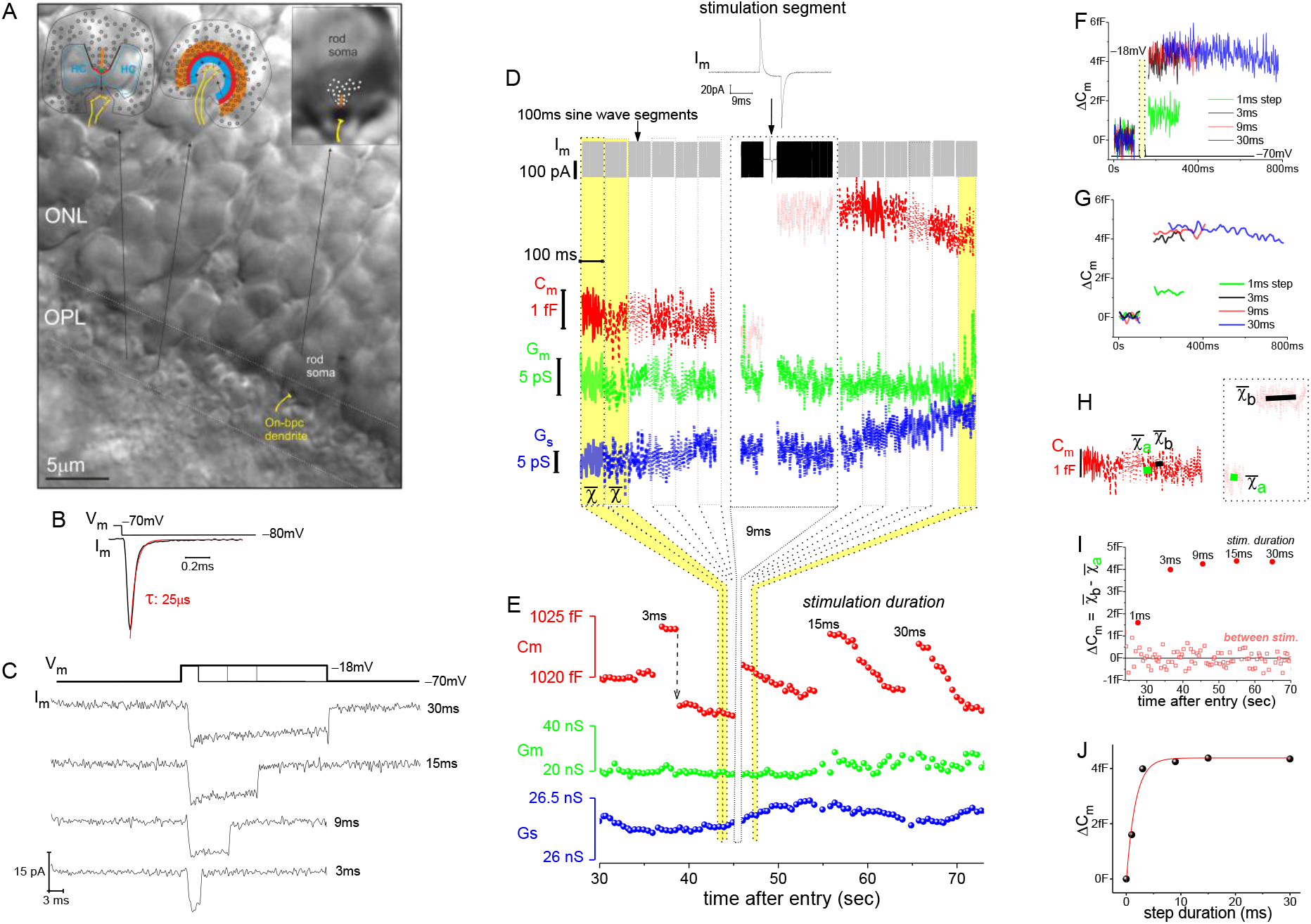
Exemplar recording from a mouse rod photoreceptor. (**A**) Image of a retinal slice centered on photoreceptor terminals in the outer plexiform layer (OPL) and somata in the outer nuclear layer (ONL). The two insets on the left and middle show rod terminals rotated 90° from one another, relative to the plane of the ribbon. The third inset shows a zoomed in view of an axonless rod with the ribbon and On-bpc dendrite added for reference. Legend: ribbon (orange); active zone (thick red line); arciform density (green diamond); ribbon flanked by synaptic ridges (thick black lines); horizontal cells (HC, in blue); and, ON-bipolar dendrite (yellow) with mGluR6 receptors (red). (**B**) The membrane current (I_m_) transient measured from a soma-ribbon in response to a brief hyperpolarizing voltage step. Current trace taken prior to compensating whole-cell membrane capacitance (C_m_). (**C**) Series of Ca^2+^ currents measured from an individual rod in response to voltage steps for the indicated durations. (**D**) Illustrates the behavior of the I_m_ and the lock-in amplifier outputs (C_m_, G_m_ and G_s_) in relation to a step depolarization. The Im presented at the top of the panel shows the stimulation segment: a 9 msec step to −18 mV. The I_m_, before and after the stimulation, was dominated by the sine wave voltage that was used to generate the lock-in outputs (membrane capacitance (C_m_), membrane conductance (G_m_) and series conductance (G_s_)). The corresponding I_Ca_ traces for this cell are presented in C. (**E**) The 100 msec sine wave segments depicted in D were binned to a mean value, and then plotted chronologically. Each stimulation produced an abrupt elevation in C_m_ (step durations indicated in figure). (**F**) Overlay of stimulated ΔC_m_ traces from E, with the baselines zeroed and traces aligned to the onset of stimulation. (**G**) Same traces as in F, but here the signals were smoothed to a corner frequency of 20 Hz to better appreciate the differences in responses (in F the f_c_ was 200 Hz). (**H**) Illustrates how ΔC_m_ was quantified over the baseline and stimulation segments. (**I**) Plots ΔC_m_ for baseline and stimulation segments, over the course of the experiment. (**J**) Summary of evoked ΔC_m_ versus step duration for this cell; step durations: 0.5, 1, 3, 9, 15 and 30 msec.

Stimulated exocytosis was followed by endocytosis of membrane, apparent as a decline in C_m_ over the extended time course (Fig. 1E). The evoked jumps in C_m_ were stationary for ~0.5 sec under the conditions used in this study (Fig. 1D-G), and then C_m_ decayed towards baseline in different ways: linearly, exponentially, and/or as abrupt downward steps (Fig. 1E). Rarely were large (> 2fF) downward steps in C_m_ observed, such as the one between the 3 and 9 msec simulations in Figure 1E (downward arrow). Given the diversity of endocytotic responses, these events were not quantified further in the current study.

### Ca_v_ channel activation was rapid and exhibited moderate inactivation

A feature that sets Ca_v_1.4 channels apart from other L-type channels is that their expression is limited to photoreceptor terminals (for review, McRory et al., 2004; Pangrsic et al., 2018). By comparison, other L-type channels, such as Ca_v_1.2 and 1.3, are expressed throughout the nervous system, and additionally in cardiac, endocrine and neuroendocrine cells, and such prevalence has led to significantly more insight into their biophysical properties (Dolphin and Lee, 2020). Therefore, some of the basic biophysical properties of mouse rod Ca_v_1.4 channels, which have not previously been described, are reported next.

The voltage-dependence of Ca^2+^-current activation was examined over a range of voltage steps, and the rods were filled with 10 mM EGTA to minimize Ca^2+^-activated Cl^−^-currents (Bader et al., 1982). The depolarization stimulated I_Ca_ showed a steep dependence on voltage (Fig. 2A), and this was quantified by measuring peak-I_Ca_ amplitude (Fig. 2B) and calcium tail-current (I_Ca_-tail) amplitude (Fig. 2C). Specifically, a depolarizing step (V_step_) to −40 mV produced a peak-I_Ca_: −0.89 ± 0.34 pA, and by −10 mV the maximal peak-I_Ca_ was reached, and averaged −14.15 ± 0.75 pA (9 cells; Fig. 2D) (liquid junction potential = 8.9 mV, and it is not subtracted from V_step_; see Methods). Fitting the peak-I_Ca_ versus V_step_ curve with a Boltzmann equation gave a half-maximal peak-I_Ca_ at a voltage of −28.7 ± 0.4 mV (see Table S3). Next, to better estimate the point when half of the available Ca_v_ channels opened (V_1/2_), a modified Boltzmann I-V equation was used, one that accounted for V_rev_ and G_max_ (see Methods). This approach gave a V_1/2_ = −23.4 ± 1.0 mV (fit presented in Fig. 2D, and Table S3). Lastly, I_Ca_-tail amplitudes were measured to determine the fraction of channels that opened at each V_step_, and this gave a half-maximal amplitude at approximately −30 mV (dashed line in Fig. 2E). However, fitting the curve with a sigmoidal equation to estimate V_1/2_ was not possible, because the I_Ca_-tail amplitudes appeared to decrease at V_step_ values positive to −10 mV (Fig. 2E). This behavior may indicate a degree of Ca_v_ channel inactivation within the 10 msec voltage step (see below). Therefore, the best estimate for half-maximal channel activation was derived from the modified Boltzmann I-V equation (Fig. 2D).

**Figure 2.**
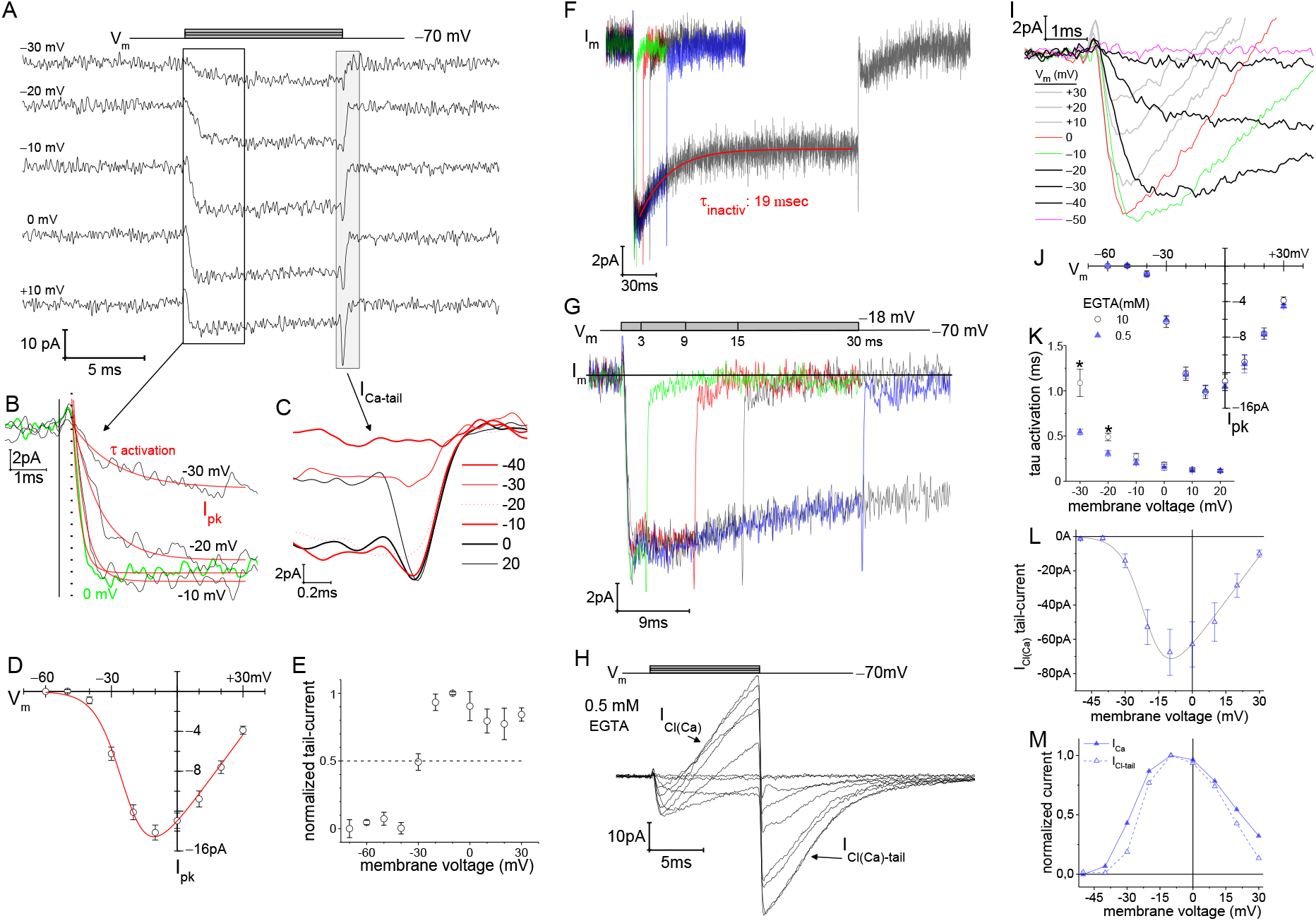
Voltage-dependence of Ca^2+^-currents and Ca^2+^-activated currents. (**A**) A family of individual I_Ca_ traces measured in response to 10 msec voltage steps, delivered at 10 mV increments, over a range of −80 to +30 mV (10 mM EGTA in the pipette). (B) Zoomed in view of I_Ca_ activation. Start of the V_step_ is indicated with the vertical solid line, and onset of I_Ca_ is indicated with vertical dashed line (dead time of ~300 μsec). An exponential function was used to fit (red lines) each I_Ca_ trace from which τ_activation_ and peak-I_Ca_ amplitude were ascertained. (**C**) Overlay of tail currents from A. (**D**) Averaged peak-I_Ca_ versus V_step_ fit with a modified Boltzmann I-V equation (V_1/2_: −24.0 ± 1.3, slope (dx): −6.2 ± 0.6 mV/e, V_rev_: +44.6 ± 2.6 mV, and G_max_: 0.30 ± 0.02 pA-mV^−1^; 9 cells; see Tables S2 and S3). (**E**) Normalized I_Ca-tail_ plotted over V_step_ (4 cells). (**F**) Overlay of average membrane currents in response to voltage steps from −70 to −18 mV for the indicated durations (averages from 7 to 13 cells). The I_Ca_ associated with the 200 msec step depolarizations (7 cells) were fit as a single exponential decay (τ = τ_inactivatιon_). (**G**) Zoomed in view of the shorter duration voltage steps highlights the rapid return of I_Ca_ to baseline (subsequent to the transient I_Ca_-tail). (**H**) A Ca^2+^-activated current emerged when the intracellular concentration of EGTA was lowered to 0.5 mM (8 cells). The outward I_Cl(Ca)_ that occurred during the step and the subsequent inward I_Cl(Ca)_-tails are designated in plot.(**I**) Expanded view of the activation portion of the current traces from C. The voltage protocol used here was the same as that in A, and the vertical line marks the start of V_step_, and the dashed vertical line indicates current onset. (**J**) Overlay of average peak-I_Ca_ plotted over V_step_, derived from experiments with 0.5 (8 cells) and 10 mM EGTA (also presented in A*ii*), are highly overlapping. (**K**) τ_activation_ versus V_step_ for low and high EGTA conditions (*: p ≤ 0.006; see Table S2). (**L**) Average I_Cl(Ca)_-tails versus V_step_ (6 cells). Boltzmann I-V fit (blue trace): V_1/2_: −20.4 ± 0.5, slope (dx): −5.6 ± 1 mV/e, V_rev_: +35.8 ± 3.8 mV and G_max_: 1.41 ± 0.44 pA-mV^−1^ (6 cells; see Table S4). (**M**) Normalized average I_Cl(Ca)_ and peak-I_Ca_ (from 0.5 EGTA data) plotted over V_step_ (error bars excluded).

Studies of Ca_v_1.4 channels in heterologous expression systems have reported they inactivate slowly (Pangrsic et al., 2018), and I_Ca_ measured from salamander rods do not exhibit inactivation (Bader et al., 1982). To assess the situation in mouse rods, prolonged steps were examined for signs of I_Ca_ inactivation. Steps from −70 mV to −18 mV, for varying durations, declined in amplitude within the first 30 msec, and after this the steady I_Ca_ was maintained (Fig. 2F). For instance, the 200 depolarizations had an initial peak-I_Ca_ = −13.2 ± 0.8 pA, and ended with a mean current I_Ca_ = −9.2 ± 1.2 pA (7 cells). This 31 % decay in I_Ca_ had a τ = 19.46 ± 0.01 msec (7 cells; Fig. 2F). For several reasons, this behavior arguably represents inactivation of Ca_v_ channels, or relaxation of facilitation, rather than contamination by other ionic currents. First, the overlay of depolarizations for the different durations shows that the membrane current approached baseline within 1 msec after repolarizing the rod (Fig. 2G), which is in accord with Ca_v_ channel deactivation (Fig. 2C). Second, the intra- and extra-cellular solutions were designed to mitigate contamination from other membrane currents (see Methods). Third, recording directly from the terminal provides optimal electrical control of the Ca_v_ channels and faster infusion (see Hagiwara et al., 2018). For these reasons, it is concluded that a fraction of the Ca_v_ channels are sensitive to inactivation, or all are partially inactivating, which to our knowledge has not been described before in photoreceptors.

### Expanding the Ca^2+^-domain triggers Ca^2+^-activated channels

By decreasing intracellular EGTA from 10 to 0.5 mM, the Ca^2+^-domain that forms about the Ca_v_ channel during depolarization is predicted to expand in size from 6 to 210 nm around the Ca_v_ channels, respectively (Neher, 1986). At ribbon synapses, 10 mM EGTA is reported to restrict the domain of elevated free Ca^2+^ to the base of the ribbon where the Ca_v_ channels reside (Moser et al., 2019), and the domain should spread up the synaptic ridges when 0.5 mM EGTA is used. When intracellular EGTA was lowered, an outward-current appeared during depolarization (presumed Cl^−^ influx), and an inward tail-current (Cl^−^ efflux) ensued upon repolarization to −70 mV (Fig. 2H). The tail-currents decayed much slower (Fig. 2H) than I_Ca_-tails (Fig. 2A-B). Since rods express the Ca^2+^-activated Cl^−^ channel TMEM-16A at their terminals (Mercer et al., 2011), and the Cl^−^ reversal potential is estimated to be −41 mV (see Methods), the Ca^2+^-activated current is presumed to be carried by Cl^−^, and referred to here as I_Cl(Ca)_ (Bader et al., 1982; Mercer et al., 2011). The magnitude of peak-I_Ca_ measured prior to the onset of the outward Cl^−^-current (measurements described in Fig. 2B; and see Methods) was virtually identical for the two EGTA buffering conditions at each V_step_ (Fig. 2J; see Table S2 and S3). In contrast, activation kinetics were influenced by changing intracellular EGTA. This was most pronounced with an intracellular concentration of 10 mM EGTA, where the tau for activation (τ_activ_) changed 6-fold when V_step_ was incrementally increased from −30 to 0 mV (1.09 ± 0.15 and 0.18 ± 0.03 msec, respectively; 9 cells; Fig. 2K). By comparison, with 0.5 mM EGTA the τ_activ_ changed 3.4-fold from −30 and 0 mV (τ_activ_ of 0.55 ± 0.03 and 0.16 ± 0.03 msec, respectively; 8 cells; Fig. 2K). The τ_activ_ values measured at 0 mV were similar with low and high EGTA, but at −30 mV the higher EGTA concentration gave a τactiv that was 2-fold slower than that measured with low EGTA (p: 0.003; see Table S2 for further kinetic descriptions). These results are interpreted as Ca^2+^-dependent facilitation of Ca_v_ channel activation kinetics, but not peak-I_Ca_ amplitude, which is similar to the behavior of inner hair cell Ca_v_1.3 channels (Goutman and Glowatzki, 2011). Such alterations in I_Ca_ activation kinetics have not been reported for PRs (for review, Pangrsic et al., 2018).

Finally, the plot of I_Cl(Ca)_-tail amplitudes versus voltage (Fig. 2L; see Table S4 for Boltzmann fits) is reminiscent of the I_Ca_-V_step_ curves (Fig. 2J). To more directly assess if the number of Cl^−^-channels activated in proportion to the number of Ca_v_ channels open, the normalized currents were plotted in parallel against V_step_ (Fig. 2M). This plot shows the I_Cl(Ca)_-tails were fully nested within peak-I_Ca_, which argues that the Ca_v_ channels serve as the primary source of Ca^2+^ for activating the Cl^−^-channels when 0.5 mM EGTA and short depolarizing steps were used. In summary, comparison of results with intracellular concentrations of 0.5 and 10 mM EGTA suggests that the Ca^2+^-activated channels are not colocalized at the base of the ribbon with the Ca_v_ channels, because 10 mM EGTA fully blocked activation of the Cl^−^-channels; but instead, they reside further up the synaptic ridges.

### The readily releasable pool of SVs is primed for ultrafast release

As the example in Figure 1 outlined, evoked ΔC_m_ reflect the number of SVs fusing. With 10 mM EGTA, the average evoked ΔC_m_ plotted against step duration showed the greatest rate of change occurred with the shortest step durations (Fig. 3A). A single exponential fit to steps from 0.5 to 9 msec was made with a τ_depletion_ = 348 μsec and an amplitude of 3.27 fF (adjusted R-square = 0.999), and extending the fit out to 30 msec gave a comparable result with a τ_depletion_ = 383 μsec and an amplitude of 3.43 fF (adjusted R-square = 0.992; 7 to 15 cells per step duration; Fig. 3A). Taking the fit to the shorter time range yields an RRP of 92 SVs. Notably, the 0.5 msec steps produced negligible changes in C_m_ (0.14 ± 0.19 fF ~4 SVs; 8 cells), while 1 msec steps released ~70 % of the RRP (ΔC_m_: 2.11 ± 0.74 fF ~60 SVs; 14 cells).

**Figure 3.**
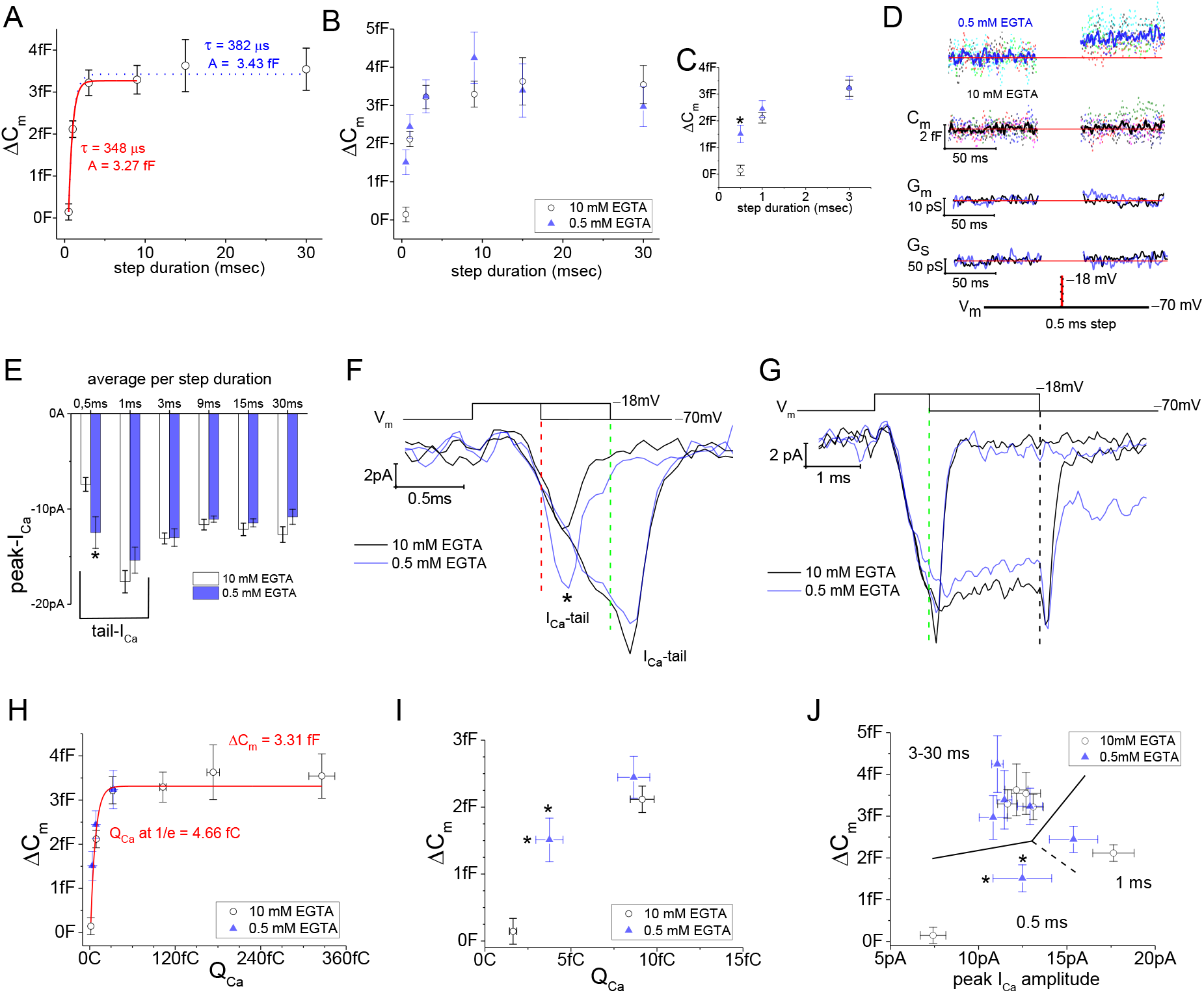
Release facilitation and ultrafast depletion of the rod’s RRP of SVs. (**A**) Average ΔC_m_ measured from rods filled with 10 mM EGTA, and stimulated with a V_step_ to −18 mV for durations: 0.5 to 30 msec. Single exponential fits to points from 0.5 - 9 msec (red curve), and from 0.5 - 30 msec (dotted curve). (**B**) Comparison of ΔC_m_ derived from experiments with 0.5 or 10 mM EGTA in the pipette; voltage step durations: 0.5 to 30 msec. (**C**) Highlights the more rapid ΔC_m_ at the singular time point of 0.5 msec when rods were loaded with 0.5 mM EGTA (*, p: 0.0016; 6 and 8 cells for 0.5 and 10 mM EGTA). (**D**) Summary of lock-in amplifier traces recorded during 0.5 msec step depolarizations, with either 0.5 or 10 mM EGTA. The C_m_ traces represent an overlay of individual recordings (dashed traces: each cell a separate color). The averaged C_m_, G_m_ and G_s_ responses are indicated by the trace font; with blue: 0.5 mM EGTA (6 cells) and black: 10 mM EGTA (8 cells). Only average responses are presented for G_m_ and G_s_. (**E**) Comparison of average peak-I_Ca_ derived from high and low intracellular EGTA. Only voltage steps for 0.5 msec were statistically different when comparing different EGTA levels (p: 0.008; see text). The 0.5 and 1 msec steps were dominated by the tail-currents, which is why their amplitudes varied from steps ≥ 3 msec. (**F** and **G**) Overlay of average current traces in response to step durations of 0.5, 1 and 3 msec (*, sign. diff.; see D). Most notable are the overlapping inward I_Ca_ traces that began after V_step_ onset, and second the moment of repolarization (vertical dashed lines) that elicited tail-currents. (**H**) Plot of ΔC_m_ over Q_Ca_. All data points from experiments with 0.5 and 10 mM EGTA were treated as one group and fit with a single exponential equation (red curve) to estimate the size of the RRP (ΔC_m_ amplitude) and the amount of Q_Ca_ to release 63 % (~1/e) of the RRP. Only steps for 0.5, 1 and 3 msec were used from the 0.5 mM EGTA dataset since an outward I_Cl(Ca)_ interfered with estimation of Q_Ca_ when step duration exceeded 3 msec. (**I**) Zoomed in view of ΔC_m_ and Q_Ca_ produced with 0.5 and 1 msec steps. * indicates a significant difference for Q_Ca_ between 0.5 mM vs. 10 mM EGTA; p: 0.04, and 6 and 9 cells, respectively. For statistical comparison of ΔC_m_ values see C and text. (**J**) Plot of ΔC_m_ over peak-I_Ca_ for step durations: 0.5 to 30 msec (dashed lines partition data points by step duration).

The change in ΔC_m_ from 3 to 30 msec was a modest 14 % increase in release (2.99 ± 0.48 vs. 3.54 ± 0.50 fF; p: 0.033, paired sample t-Test; 7 cells). To derive a steady release rate, 200 msec depolarizations were made later in the recordings, outside the series of short duration steps. After accounting for potential rundown in release (see Fig. S1A-F), the 200 msec depolarizations are estimated to yield 5.31 ± 1.25 fF (Fig. S1D; 7 cells). Subtracting this value from the initial exponential release phase (the RRP) gives a ΔC_m_ of 2.04 fF (= 5.31 fF – 3.27 fF) within 200 msec, which equates to a continual release rate of 10.2 fF-s^−1^. In total, the results provide clear evidence for an ultrafast pool of 92 SVs with a τ_depletion_ < 0.4 msec, and subsequent release under steady Ca^2+^ entry proceeded at a rate of 288 SVs-s^−1^.

### Lowering the intracellular Ca^2+^-buffer expedited release onset

When rods, filled with 0.5 mM EGTA, were depolarized for 0.5 msec to −18 mV, the ΔC_m_ was over 10-fold larger than the responses measured with 10 mM EGTA (1.51 ± 0.36 vs. 0.14 ± 0.19 fF; p: 0.006; n: 6 and 8 cells; Fig. 3B and C). This is also illustrated with a comparison of the individual and average C_m_, G_m_, and G_s_ traces from the recordings with low and high EGTA (Fig. 3D). In contrast to the initial distinction in ΔC_m_, over the range of 1 to 30 msec steps there was a high degree of overlap between the two datasets (Fig. 3B), indicating the RPP of SVs was not altered by changing the concentration of intracellular EGTA. To understand what expedited the onset of vesicle fusion, the kinetics of channel activation were considered for the two Ca^2+^-buffering conditions.

Since a delay of ~300 μsec existed between the start of depolarization and I_Ca_ onset (Fig. 2B and 2I), there was only ~200 μsec for I_Ca_ to develop before the 0.5 msec step ended and the V_m_ repolarized; hence, the 0.5 msec steps created what were essentially tail-currents. Given that τ_activ_ at −20 mV for low and high EGTA were 310 versus 489 μsec, respectively (Table S2; Fig. 2K), it is expected that more channels opened with less Ca^2+^ -buffering. The results show that the average I_Ca_-tail resulting from 0.5 msec steps to −18 mV were approximately two-fold larger when less EGTA was used (−7.4 ± 0.7 versus −12.5 ± 1.7 pA; p: 0.008 from 9 and 6 cells; Fig. 3E and F). In contrast, the 1 msec steps formed comparable I_Ca_-tail under the two intracellular EGTA conditions (Fig. 3E-G), because the step duration was > τ_activ_. Finally, the current responses to longer depolarizations (≥ 3 msec) were separable from the tail-currents and quantified by measuring the initial peak-I_Ca_ (as illustrated in Fig. 2B). These longer steps gave similar peak-I_Ca_ that ranged from −11 to −13 pA in both low and high Ca^2+^-buffering (Fig. 3E). This result agrees with the data derived from the 10 msec steps that were used to construct I-V curves (Fig. 2J), specifically: peak-I_Ca_ was the same under conditions of low and high EGTA even though activation times were different.

Evaluating ΔC_m_ as a function of the Ca^2+^ influx, rather than time, again emphasizes the finite size of the RRP. Since Q_Ca_ is the integral of a I_Ca_ that inactivated moderately over time (Fig. 2F), the relationship between ΔC_m_ and Q_Ca_ resembled that observed for ΔC_m_ versus step duration (Fig. 3H). The combined ΔC_m_ and Q_Ca_ data from low and high EGTA experiments (11 and 15 cells, respectively) formed a continual exponential curve with a ΔC_m_ amplitude = 3.31 fF (RRP ~ 94 SVs; Fig. 3H). A Q_Ca_ at 1/e reached 4.66 fC, which equates to 12,263 Ca^2+^ ions (1 Ca^2+^ ion = 3.8×10^−19^C) that are needed to empty 63 % of the RRP of SVs (Fig. 3H). The second comparison made here is ΔC_m_ versus I_Ca_ amplitude. Plotted in Figure 3J is a combination of I_Ca_-tail (from 0.5 and 1 msec steps) and peak-I_Ca_ measured at the onset of depolarization (steps ≥ 3 msec). The first observation is that steps ≥ 3 msec in duration produced a group of ΔC_m_ values between 3 to 4 fF, i.e. depleted the RRP, at a peak-IC_a_ of approximately −12 pA (Fig. 3J). In contrast, the four IC_a_-tails stimulated with 0.5 and 1 msec steps produced a range of ΔC_m_ (0.2 to 2 fF) and I_Ca_ amplitudes (−7.5 to −18 pA) that approximated a linear relationship (Fig. 3J). In summary, a smooth exponential depletion of the RRP is apparent when ΔC_m_ is analyzed as a function of Q_Ca_ or the duration of depolarization.

Electron microscopy (EM) studies of mouse and cat rod ribbons have described some 60 to 120 SVs docked at the base of the ribbon (Rao-Mirotznik et al., 1995; Zampighi et al., 2011; Cooper et al., 2012; Grabner et al., 2015), which correlates well with the physiological findings presented here. An additional 300 SVs were reported near the plasma membrane, along the synaptic ridges, which run vertically from the base of the ribbon (Zampighi et al., 2011; for review, Moser et al., 2019). These SVs are not ribbon-tethered, nor are they necessarily tightly associated with the plasma membrane, i.e. docked. The evoked exocytosis responses presented here can be interpreted as release originating from the base of the ribbon since 1) the RRP is close to the number of SVs estimated to be at the base of the ribbon, and 2) expanding the domain from < 10 nm (10 mM EGTA) to ~200 nm (0.5 mM EGTA) did not recruit more SVs (Fig. 3B).

### Ribbonless rods support only marginal evoked exocytosis

An earlier electron microscopy study that introduced RIBEYE-ko mice, which lack synaptic ribbons in PRs, showed that ribbonless rods had only 40 % of wild type (wt) SVs at their active zones (Maxeiner et al., 2016). The study did not measure release from rods; therefore, we tested whether the loss of SVs would show up as a correlated reduction in the size of the RRP. The average ΔC_m_ following 1 and 3 msec steps delivered to ribbonless rods filled with 10 mM EGTA amounted to 0.21 ± 0.20 and 0.68 ± 0.34 fF, respectively (Fig. 4B; 7 and 8 cells), which were 10- and 5-fold smaller than wt ΔC_m_ evoked with 1 and 3 msec steps: 2.12 ± 0.20 and 3.22 ± 0.31 fF (Fig. 3A), respectively (p-values < 0.0001 for wt vs. ko; 14 and 15 cells). Responses from ribbonless rods elicited with longer duration steps did not exceed the ΔC_m_ elicited by 3 msec steps (Fig. 4A). Fitting ΔC_m_ versus step duration with a single exponential equation gave a τ_depletion_ = 560 μsec and a ΔC_m_ amplitude = 0.84 fF (RRP ~ 24 SVs; Fig. 8 cells), i.e. only 24 % the size of that measured in wt rods (ΔC_m_: 3.43 fF).

**Figure 4.**
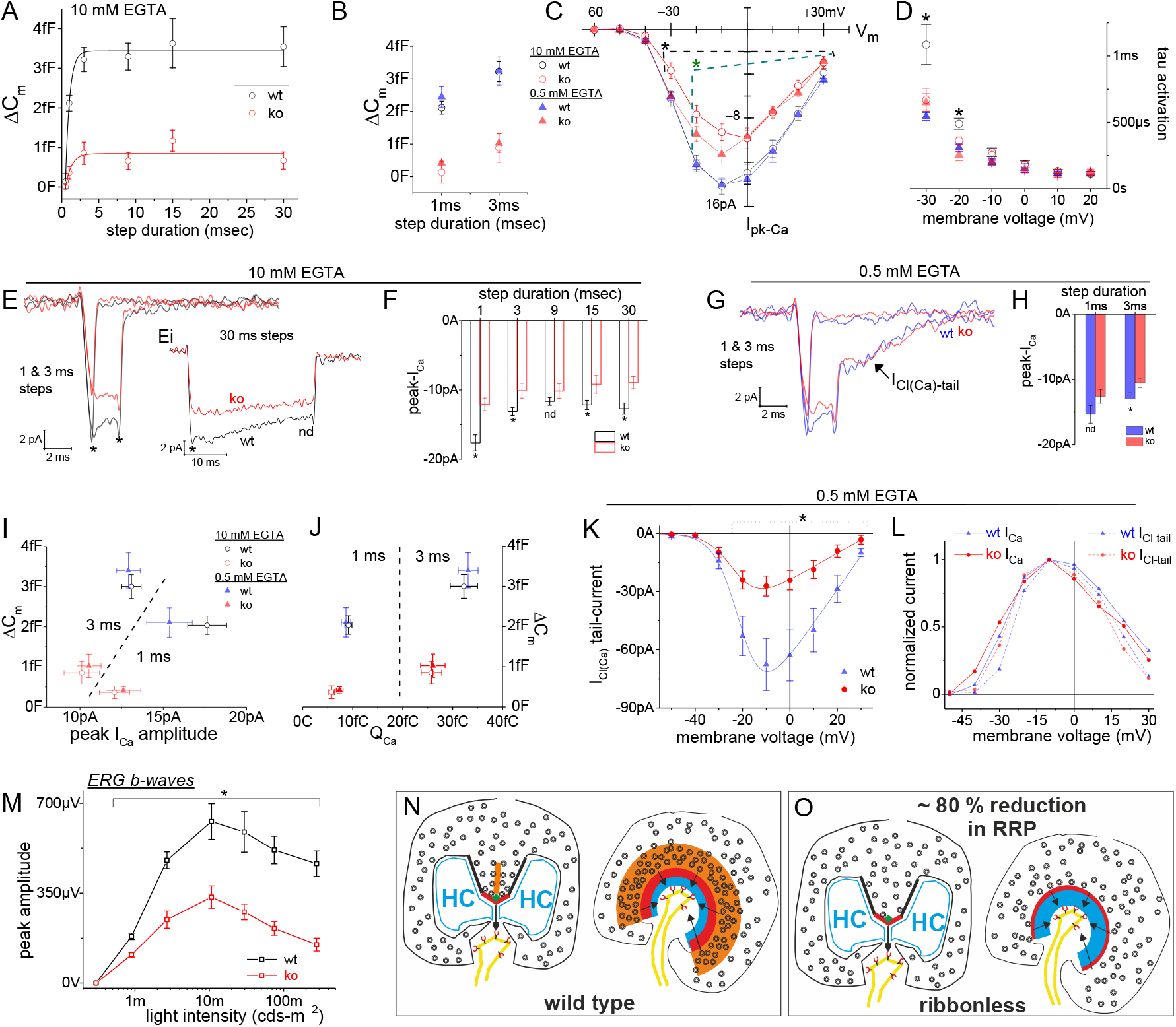
Ribbonless rods lack Ca_v_ channel facilitation, form a modest RRP and lack normal light responses. (**A**) Estimation of RRP size and depletion kinetics for RIBEYE-ko rods (10 mM EGTA in the pipette; step durations ranged from 1 to 30 msec). Wild type responses overlaid for comparison, and their evoked ΔC_m_ were significantly larger than ko values at every step duration (see text for description of fits). (**B**) Comparison of all responses that were evoked with 1 and 3 msec steps, with either 0.5 or 10 mM EGTA in the pipette, and for wt and ko rods. All ko responses were significantly smaller than wt values, see text. (**C**) The peak-I_Ca_ plotted over V_step_. Ribbonless rods had significantly smaller peak-I_Ca_ at the indicated V_steps_ (*, p-values < 0.05), with comparisons made between wt vs. ko rods for 0.5 mM EGTA (green dashed line) and 10 mM EGTA (dashed grey line). See Fig. 2A for a description of voltage step protocols and analysis. (**D**) I_Ca_ activation kinetics were significantly slower for wt rods filled with 10 mM EGTA (* indicates sig. diff.; see Table S2 for details). (**E** and **F**) Averaged I_Ca_ traces illustrates the differences between genotype were pronounced shortly after the onset of depolarization (peak-I_Ca_) and at the end of short duration steps (*, at onset and tail-currents). (**Ei**) Wild type rods filled with 10 mM EGTA showed a faster rate of I_Ca_ inactivation than ko rods. At the end of the 30 msec steps, the traces were not statistically different (nd, p > 0.3). (Note: 1msec steps were tail-currents, and therefore larger in amplitude than peak-I_Ca_.). (**G** and **H**) Comparison of genotypes with 0.5 mM EGTA. Statistical comparisons showed no difference for 1 msec steps (nd, p: 0.13; 9 and 11 cells each), while the 3 msec steps were significantly larger in wt (*, p: 0.05; 9 and 12 cells each). (**I** and **J**) All ΔC_m_ in response to 1 and 3 msec steps plotted over peak-I_Ca_, and Q_Ca_. (**K**) Average I_Cl(Ca)_-tails versus V_step_. For the Boltzmann I-V fits see Table S4. (**L**) Normalized average I_Cl(Ca)_-tails and peak-I_Ca_ plotted over V_step_ for wt and ko rods (error bars excluded). (**M**) Summary of dark-adapted erg b-waves measured from wt and ko mice (*, p < 0.04; 4 ko and 3 wt mice). (**N**) Cartoon of a wt rod and (**O**) a ribbonless rod.

To further investigate if the reduction in exocytosis measured from ribbonless rods reflected a reduction in the number of SVs available for release or impairment of coupling of Ca^2+^ influx to SVs, as concluded to occur at ribbonless retinal bipolar cells (Maxeiner et al., 2016), the concentration of intracellular EGTA was lowered. Ribbonless rods filled with 0.5 mM EGTA and given 1 and 3 msec stimulations produced ΔC_m_ that averaged 0.70 ± 0.16 and 0.86 ± 0.67 fF, respectively (Fig. 4B; 12 cells for each group). In contrast, wt rods filled with 0.5 mM EGTA given 1 and 3 msec stimulations averaged ΔC_m_ of 2.44 ± 0.31 and 3.24 ± 0.43 fF respectively, (Fig. 4B; 11 cells in each group); thus, ribbonless rods generated evoked ΔC_m_ that were < 30 % the size of wt responses (p-values for wt vs. ko, 1 and 3 msec: 0.00016 and 0.00023). Overall, the results from the different Ca^2+^-buffering conditions indicate that the RRP formed by ribbonless rods was only ~24 SVs, which is approximately a quarter the size of the RRP of wt rods. Interestingly, the different EGTA concentrations did not significantly influence RRP size within either genotype, nor were the kinetics of depletion altered in ko mice. Thus, these results suggest that the coupling of Ca^2+^ influx to exocytosis was not noticeably different between genotypes, but rather the number of release ready SVs appears to be the primary distinction between ribbonless and wt rods.

### Deleting the ribbon alters Ca_v_ channel properties

Previous work had indicated the density of Ca_v_1.4 staining was altered in ribbonless rods (Maxeiner et al., 2016; Dembla et al., 2020). More to the point, the ribbon-shaped profile that Ca_v_ channels adhere to in control rods were reduced in length by 50 % in ribbonless rod terminals; however, protein levels of the α1F subunit of Ca_v_1.4 were similar in Western blots of wt and ko retina (Maxeiner et al., 2016). To test if these changes affect the behavior of Ca_v_1.4 channels, a comparison of Ca^2+^ currents from wt and ribbonless rods was made. Results derived from 10 msec voltage steps showed a significant overall reduction in the ribbonless rods peak-I_Ca_ across V_step_’s from −30 to +30 mV (Fig. 4C; and see Tables S2 and S3). Specifically, with 10 mM EGTA in the pipette, the peak-I_Ca_ amplitude was approximately 35 % smaller in ko rods than in wt rods (p < 0.001 or smaller, depending on V_step_; 9 wt and 5 ko cells), and when 0.5 mM EGTA was used the amplitudes were 20 % smaller than in wt controls (p < 0.05 or less; 8 wt and 7 ko cells; Fig. 4C, and Table S3). Additional biophysical values were derived from Boltzmann fits to the I-V curves. This analysis shows that ko rods had a maximal conductance ~ 34 % smaller than wt values, and this was true for experiments performed with high and low intracellular EGTA concentrations (p: 0.024 and 0.029, respectively; Table S3). In addition, with 10 mM EGTA in the pipette V_1/2_ for peak-I_Ca_ amplitude measured from ribbonless rods was shifted +1.4 mV from wt values (though the level of significance depended on the specific type of Boltzmann fit; Fig. 4C and see Table S3). In contrast, 0.5 mM EGTA tended to shift ribbonless rods V_1/2_ in the opposite direction by −1.7 mV from wt values, but again the difference was not significant (p ~ 0.17; 8 and 7 cells; Fig. 4C and see Table S3). The final comparison made from the voltage steps are current activation kinetics, which were similar under all conditions, with one exception: Ca_v_ channels in the wt rods filled with 10 mM EGTA were significantly slower to open (Fig. 4D).

The reduction in ribbonless rod peak-I_Ca_ was also observed in recordings that depolarized rods to −18 mV for different durations with 10 mM EGTA in the intracellular solution (Fig. 4E and 4F); however, over time the difference faded as Ca_v_ channels in wt rods inactivated more rapidly. Specifically, 30 msec steps had an initial peak-I_Ca_ that was larger in wt rods (−13.3 ± 0.43 pA vs. −8.7 ± 0.8 pA, p: 0.0002; 7 per genotype), but by the end of the 30 msec step the wt and ribbonless I_Ca_ were no longer statistically different (−8.7 ± 0.8 pA vs. −7.4 ± 1.1 pA, p: 0.35; 7 cells each; Fig 4Ei). In the case of recordings with 0.5 mM EGTA in the pipette, the 1 and 3 msec depolarizations tended to produce smaller I_Ca_ amplitudes in the ribbonless rods (Fig. 4G, and see 4H for statistical comparisons). In total, Ca_v_ channel behavior in ribbonless rods filled with low or high EGTA were distinguished from their wt counter parts in one or more of the following ways: they had a lower I_Ca_ density at the onset of depolarization, showed a shift in V_1/2_ and they exhibited less current inactivation (or less facilitation at onset). The influence of peak-I_Ca_ and Q_Ca_ on evoked ΔC_m_ are summarized for 1 and 3 msec steps, with low and high intracellular EGTA, and for both genotypes (Fig. 4I and J). In the end, the ribbon was the most significant determinant of the magnitude of the evoked ΔC_m_.

### Alterations in ribbonless rods Ca^2+^-activated currents

The I_Cl(Ca)_-tails currents obtained in 0.5 mM EGTA from wt rods (Fig. 2C), followed the behavior of Ca_v_ channel activation (Fig. 2L and M), and same can be said for I_Cl(Ca)_-tail currents measured from ko rods (Fig. 4K). However, ribbonless rods produced significantly smaller I_Cl(Ca)_-tails amplitudes, which were only 40 % the size of wt currents (Fig. 4K; Table S4). To further evaluate the voltage-dependence of current activation, Boltzmann fits were made to the I_Cl(Ca)_-tails versus V_step_ curves. On average, the maximal conductances measured from ribbonless rods were 50 % smaller than currents recorded from wt rods (G_max_: 0.70 ± 0.13 versus 1.41 ± 0.44 pA-mV^−1^), though the difference was not statistically significant (p: 0.17; see Table S4). This parallels the observation that the peak-I_Ca_ amplitude was significantly larger in wt than ko rods (32 % larger; Table S3). Next, the voltage for half-maximal I_Cl(Ca)_ was reached at more negative membrane voltages than wt rods (V_1/2_ for I_Cl(Ca)_, wt vs. ko: −20.4 ± 0.5 vs. −22.8 ± 0.6 mV, respectively; p: 0.016. Table S4). This correlates with the tendency for peak-I_Ca_ amplitude measured from ribbonless rods to have a negatively shifted V_1/2_ when 0.5 mM EGTA was in the pipette (peak-I_Ca_ V_1/2_ values, wt vs. ko: −23.2 ± 0.9 vs. −25.3 ± 1.1 mV, respectively; p: 0.17, Table S3). Overall, the Ca^2+^-activated currents were fully nested within the peak-I_Ca_, indicating their dependence on Ca^2+^, and there was a tendency for ribbonless rods I_Ca_-V and I_Cl(Ca)_-V curves to shift to more hyperpolarized voltages relative to wt curves. This is illustrated in plots of normalized current amplitude versus membrane voltage for the ribbonless and wt rods (Fig. 4L).

### Light responses in ribbonless mice are greatly reduced

The results so far show a stark reduction in the RRP of SVs. In the intact animal, this deficit is expected to preferentially impact rod signaling in the dark, a period when the rate of glutamate release is the highest. Normally, presenting a dim light flash to a dark-adapted retina will cause a momentary pause in glutamate release from rods, which in turn causes a depolarization (dis-inhibition) of postsynaptic rod bipolar cells. The magnitude of the light response, as assessed with electroretinogram (erg) recordings, will reflect the number of rod to rod-bipolar synapses that detect a transient drop in glutamate. If RIBEYE-ko animals are unable to keep glutamate levels high enough to inhibit normal numbers of rod bipolar cells in the dark, then their light responses should be smaller than their wt counterpart. To test this hypothesis, ergs were performed on dark adapted mice, under scotopic test conditions. The a-wave component of an erg represents phototransduction in the outer segments, which hyperpolarizes the rods. Wild type and RIBEYE-ko mice had nearly identical a-wave amplitudes across the range of flash intensities (Fig. S2C), which suggests phototransduction in ribbonless rods was not altered. In contrast, the b-wave, which originates from depolarizing rod bipolar cells, measured from RIBEYE-ko animals were much smaller, reaching only 52 to 38 % of the peak amplitude of wt responses. This difference was significant over two decades of flash intensities and widened with each incremental increase in flash intensity (p-values ranging from 0.04 to 0.005; Fig. 4M). Further description of erg kinetics involved comparing the rate of rise for the a- and b-waves. The b-waves recorded from wt mice rose on average two-fold faster than b-waves measured from RIBEYE-ko mice (p-values ranging from 0.04 to 0.009; Fig. S2F). Finally, there was no difference between genotypes when it came to the rate of rise for the a-waves (Fig. S2G). These findings arguably show that ribbonless rods maintained an anemic glutamate gradient across their synapses, which was unable to inhibit a large cohort of postsynaptic rod bipolars; thus, a pause in release was not noticed.

## Discussion

The idealized functionality of the rod’s synaptic architecture is to equalize release sites (Rao-Mirotznik et al., 1998). At the cat rod synapse the shortest and longest distances between the ribbon and dendrite averaged 130 and 640 nm, respectively (Rao-Mirotznik et al., 1995). These distances are estimated to create peak synaptic glutamate concentrations that vary by 100-fold, which is arguably too much variability for an iontotropic GluR synapse to utilize quantal signaling (Rao-Mirotznik et al., 1998). To counter this, ON-bipolars express high affinity mGluR6 receptors that have an EC50 for glutamate that is ~10 μM, which are much higher affinity than iGluRs (see references within: Rao-Mirotznik et al., 1998; DeVries et al., 2006); furthermore, the mGluR6 cascade is prone to saturate (for review, Field and Sampath, 2017). These features may enable release events from different distances to have equal effect on the postsynapse. In the current study, the results show that a rod ribbon emptied its RRP in a single kinetic phase with a τ_depletion_ of ~0.4 msec (Fig. 3A), and the size of the RRP was equally large (~92 SVs) when rods were filled with 0.5 or 10 mM EGTA (Fig. 3B). The ultrafast rate of depletion strongly suggests that only primed SVs were released (Neher and Brose, 2018). Since the RRP fused as a single exponential phase and RRP size was not altered by changing intracellular EGTA levels, we suggest that the SVs are uniformly primed and within nanometers from Ca_v_s channels.

The presumed stretch of release sites at the base of a mouse rod ribbon has a contour length of 1.6 μm (measured from RIBEYE profiles; Grabner et al., 2015; 1.7 μm EM-FIB, Hagiwara et al., 2018). Two separate analyses of EM tomograms estimated ~60 (Zampighi et al., 2011) and ~77 (Cooper et al., 2012) docked SVs at the ribbon base. These values are slightly less than the maximal, calculated number of 86 SVs that can be packed in this region (Grabner et al., 2015) (see schematic in Fig. 4N). The physiological measurements indicate ~92 SVs in the RRP (Fig. 3B), which is 6 to 20 % greater than the calculated and measured number of SVs at the ribbon’s base, respectively. The correlation between the anatomy and evoked responses supports the notion that the rod’s large ribbon AZ spans the length of the crescent-shaped ribbon. The synaptic, mathematical models outlined above conclude that a minimal steady release rate ≥ 40 SVs-s^−1^ is needed to keep the rod ON-bipolar pathway continually activated (Rao-Mirotznik et al., 1998; Hasegawa et al., 2006). We find that the rod ribbon release rate is 288 SV-s^−1^ during a sustained, strong depolarization to −18 mV. Assuming each of the 92 SV release sites participate equally during sustained release, then each site refills at 3 Hz. At a more physiological membrane voltage close to V_1/2_ there should be sufficient Ca^2+^ influx to sustain > 40 SV-s^−1^, which translates to a turnover rate of ~0.40 Hz per release site.

Comparing results from mouse rods to studies carried out on isolated Mb1 bipolars highlights differences in the timing of SV fusion. A subpopulation of Mb1 SVs fuse with a τ_ultrafast_ ~0.5 msec (Heidelberger et al., 1994; Mennerick and Matthews, 1996; Burrone et al., 2002; Palmer et al., 2003), which are rate limited by Ca_v_ channel activation kinetics (τ ~0.6 msec at −10 mV, Mennerick and Matthews, 1998), and their release is unimpeded by elevated intracellular Ca^2+^-buffering (5 mM EGTA) (Mennerick and Matthews, 1996). An additional population of SVs are considered to reside at greater distances from Ca_v_ channels, because they only enter the RRP when intracellular Ca^2+^-buffering is reduced (0.1 mM EGTA) (Burrone et al., 2002). The heterogeneity in the Mb1’s primed SV release kinetics (Grabner and Zenisek, 2013) is not observed in the mouse rod experiments. Though the mouse rod RRP of SVs are tightly and uniformly coupled to Ca_v_ channels, we find that facilitation of Ca_v_ channel activation kinetics occurs on a timescale that accelerates the rate of fusion. In response to 0.5 msec voltage steps, the RRP was depleted by 50 % with 0.5 mM EGTA, but only 4 % of the RRP emptied with 10 mM EGTA (Fig. 3Bi). The faster release onset is attributed to the acceleration of Ca_v_ channel activation kinetics in 0.5 mM EGTA (Fig. 2E), which enhanced Q_Ca_ selectively at 0.5 msec (Fig. 3D and 3H). Interestingly, Burrone et al. (Burrone et al., 2002) showed something very similar with respect to release, specifically a tail-current released 50 % of the RRP when the Mb1 bipolars were loaded with 0.1 mM EGTA, and with higher Ca^2+^ buffering (endogenous) the tail-current released only 5 % of the RRP. They attributed the enhanced rate of release to the expansion of the Ca^2+^-domain, but differences in I_Ca_ were not reported.

Auditory hair cells also support a robust form of facilitation when they are pre-conditioned with elevated, free intracellular Ca^2+^, which is achieved by pre-depolarizing the cell and/or lowering intracellular Ca^2+^-buffering. Release facilitation was characterized by shorter onset latencies and higher release synchrony (Cho et al., 2011; Goutman and Glowatzki, 2011). Notably, rat inner hair cells (IHCs) showed facilitation of I_Ca_ onset, but without a change in steady-I_Ca_ amplitude (Goutman and Glowatzki, 2011), similar to what we report for rods, while frog HCs did not exhibit a change in I_Ca_ onset (Cho et al., 2011). These studies show that mammalian IHCs (Goutman and Glowatzki, 2011) and rods accelerate Ca_v_ channel opening (this study), possibly via elevating basal Ca^2+^, which expedites release. This suggests that release is limited by Ca_v_ channel activation kinetics. In contrast, frog HCs (Cho et al., 2011) and Mb1 bipolars (Burrone et al., 2002) accelerate release through a distinct process that may involve a Ca^2+^-dependent priming step or differences in the spatial coupling between Ca_v_ channels and SVs, respectively.

To further test the ribbon-centric release model, evoked SV turnover was studied in ribbonless rods. Here the RRP was whittled down to 24 SVs, compared to 92 SVs in wt rods (Fig. 4A). Since the extent of release was nearly identical with high and low intracellular EGTA concentrations, the RRP formed by ribbonless rods is tightly coupled to the Ca_v_ channels as in wt rods (Fig. 4B). The ribbonless RRP may originate from the AZ territory where the base of the ribbon and arciform density are normally located in wt rods (depicted schematically in Fig. 4N and 4O). The density of SVs in this region amounted to 40 % of the normal number of SVs present in wt rods (Maxeiner et al., 2016), and the length of the horseshoe-shaped Ca_v_1.4/RIM2 profiles were reduced by 50 % (Dembla et al., 2020). Thus, a 50 % loss of AZ area, and only 40 % the normal density of SVs, can account for the deficiencies in the size of the RRP. Another potential source of SVs are the synaptic ridges that run vertically some ~200 nm from the base of the ribbon, and amount to ~300 SVs in wt rods (Zampighi et al., 2011). Their contribution to evoked responses is likely minor in wt rods, because changing Ca^2+^-buffering did not alter pool size, which argues against a distant extra-ribbon source of SVs (see schematics in Fig. 4N and O). Furthermore, rather than supporting release, the ridges are associated with endocytotic structures and the related molecular markers for endocytosis (Wahl et al., 2013; Fuchs et al., 2014; for review Moser et al., 2019). Taken together, it is possible that wt rods form an RRP from ~70 SV-release sites that are ribbon-dependent, and ~20 additional SVs are within the AZ, but independent of the ribbon (Fig. 4O). In contrast, experiments on salamander rods that acutely disrupted ribbon AZs by photo-damaging them, showed that ΔC_m_ following a 25 msec depolarization was reduced by 50 % after photo-damage, but longer depolarizations to 200 msec were similar to control (non-damaged) responses (Chen et al., 2013). This was interpreted as salamander rods forming a large extra-ribbon RRP of SVs, which fits nicely with the observations that they have multiple kinetic phases of release (Kreft et al., 2003; Rabl et al., 2005). In addition, salamander rods express 5 to 7 ribbons per terminal, which may create more release heterogeneity between ribbons. Mouse rods are comparitively less complex, expressing only a single ribbon that empties in a single ultrafast step.

In this study we also observed that wt peak-I_Ca_ decayed by a third with a τ of 19 msec, and this occurred with 10 mM EGTA in the pipette. Upon repolarization, the I_Ca_ traces approached baseline in < 1 msec (Fig. 2C, 2G, 3F and 4Ei), which fits the description of Ca_v_ channel deactivation rather than involvement of Ca^2+^-activated currents. This degree of I_Ca_ inactivation was not observed in salamander rods (Bader et al., 1982), and measurements of I_Ca_ from porcine (Cia et al., 2005) and ground squirrel rods (Li et al., 2010) did not report I_Ca_ inactivation. However, the charge carrier used in one study was 2 mM Ba^2+^ (porcine rods) and in the other study 10 mM BAPTA was used to buffer Ca^2+^ (squirrel rods), which eliminate a likely candicate mechanism: Ca^2+^-dependent inactivation (CDI). There is growing evidence that heterologously expressed human Ca_v_1.4 in HEK cells can produce CDI (Haeseleer et al., 2016) that is activated by PKA, which causes an increase in peak-I_Ca_ amplitude (enhanced open probability) that inactivates over time (Sang et al., 2016). Second, raising the recording temperature from 23°C to 37°C has been reported to strongly increase peak-I_Ca_ amplitude (by 3-fold) and accelerate voltage-dependent inactivation (VDI) of Ca_v_1.4 currents in HEK cells (by 50-fold) (Peloquin et al., 2008, with 20 mM BaCl2). We point out that previous I_Ca_ measurements from mouse rods at 28°C (Grabner et al., 2015) generated ~50 % less peak-I_Ca_ amplitude than what was observed in the current study (~31°C) and another recent study performed at a higher temperature (Wang et al., 2017; Hagiwara et al., 2018). Future studies will need to further assess whether CDI and/or VDI are involved, and more directly test if inactivation is temperature-dependent.

Both of the Ca_v_ channel gating phenomena observed in wt rods were eliminated in ribbonless rods. I_Ca_ activation kinetics measured with 10 mM EGTA in the pipette showed slower activation kinetics in wt than ribbonless rods (Fig. 4D and Table S2), and inactivation was only observed in wt rods (Fig. 2F and 4Ei). Given the steady-I_Ca_ amplitude measured from wt and ko rods were similar, there were arguably comparable numbers of open channels at ~30 msec; however, ribbonless rods appear to lack a transient enhancement in I_Ca_ (channel open probability). Since normal levels of Ca_v_1.4 (α1F-subunit) protein (as well as RIM2α) are expressed in retina from RIBEYE-ko mice (Maxeiner et al., 2016), it is not surprising that the different genotypes have comparable steady-I_Ca_ amplitudes. What has been disrupted in ribbonless rods are the horseshoe-shaped AZ profiles for Ca_v_1.4 (α1F) and RIM2α, which are 50 % shorter in length than wt AZs (Maxeiner et al., 2016; Dembla et al., 2020). Since loss of interactions between Ca_v_1.4 and AZ interacting partner RIM2α reduced rod peak-I_Ca_ by 80 % (Grabner et al., 2015), it’s likely that removing the ribbon impaired these facilitatory interactions.

Our study provides new insight into the biophysics of SV fusion at mammalian rod ribbons. The results demonstrate that the rod ribbon creates multiple release sites with similar release probability in response to a strong stimulation. This is driven by Ca_v_ channels that activate at ultrafast rates. The coupling between Ca_v_ channels and SVs is on a nano-scale, and no signs of heterogeneity in spatial coupling were apparent. Instead release heterogeneity arose for Ca_v_ channel facilitation, which was specific to the timing of release onset. These features were dependent on the synaptic ribbon, as ribbonless rods lacked Ca_v_ channel facilitation and the RRP was greatly scaled down. Future studies should attempt to determine the functional stoichiometry of a release site, starting with the following questions: how many Ca_v_ channels open to trigger a SV(s) to fuse, what mechanisms impact release probability (i.e., PKA), and is release site uniformity maintained when weaker depolarizations are used. Better insight into these matters will further our understanding of how rods convert depolarizations into synaptic signals, and ultimately how the mammalian rod pathway encodes object motion and position (Tikidji-Hamburyan et al., 2017; Field et al., 2019).

## Materials and Methods

### Animal handling

Animals were handled in accord with institutional and national animal care guidelines. The *ribeye* knockout mice (Maxeiner et al., 2016) were a kind gift from Frank Schmitz and Stefan Maxeiner (University of Saarland). They were maintained on a C57BL6/J background and crossed as *ribeye*^−/+^ Wild type and ko littermates, male or female, and between 3 and 6 months of age were used for experiments during the daylight phase of the day/night cycle.

### Electrophysiology

Retinae were dissected at an ambient temperature of 18-20°C and then submersed in mouse extracellular solution (MES) with a low Ca^2+^ concentration that had the following composition (in mM): 135 NaCl, 2.5 KCl, 0.5 CaCl_2_, 1 MgCl_2_, 10 glucose, 15 HEPES, and pH adjusted to 7.35 with NaOH and an osmolarity of 295 mOsm. Dissected portions of retina were absorbed onto pieces of nitrocellulose membrane mounted onto glass with the vitreal side the retina contacting the membrane. The sclera and pigment epithelium were from the exposed surface of retina and then ~200 μm thick slices were made with a custom built tissue chopper. Immediately after slicing, the retinal sections (attached to the nitrocellulose membrane) were transferred to the recording chamber and arranged to be viewed in vertical cross section to optimize resolution of the OPL. Slices were washed continually with low Ca^2+^ MES for approximately 5 minutes as they equilibrated to an ambient temperature of 30° to 32°C, and then the Ca^2+^ was increased to 2 mM.

The intra- and extra-cellular recording solutions have been described previously (Grabner et al., 2015), and a few modifications were made to improve the I_Ca_ measurements. To further block a delayed rectifier, outward K^+^-current (Cia et al., 2005), the concentration of TEA was increased to 20 mM in the intrac- and to 35 mM in the extracellular solutions. In addition, Cs^+^ replaced intracellular K^+^. Next, a glutamate transporter Cl^−^-current was previously blocked with high concentrations of DL-TBOA, 350 μM (Grabner et al., 2016), which is a non-selective EEAT blocker. In the current study we used TFB-TBOA, which is a high affinity blocker for EEAT1-3, at a concentration of 3 μM (TOCRIS). It showed better stability over time, and far greater potency than 350 μM DL-TBOA (Grabner et al., 2016). The terminals have an I_h_ current (Hagiwara et al., 2018) that was blocked by adding 5 mM CsCl to the extracellular solution (Bader et al., 1982). Finally, extracellular HEPES was elevated to 15 mM to block inhibitory proton feedback onto Ca_v_ channels (find reference in: Grabner et al., 2015). In the end the extracellular recording solution had the following reagents (mM): 105 NaCl, 2.5 KCl, 35 TEA-Cl, 5 CsCl, 2 CaCl2, 1 MgCl2, *0.003* TFB-TBOA, 15 HEPES, and pH adjusted to 7.35 with NaOH, and a final osmolarity between 290 to 295 mOsm. The intracellular solution with 10 mM EGTA consisted of the following reagents (mM): 105 CsCH3SO4, 20 TEA-Cl, 1 MgCl2, 5 MgATP, 0.2 NaGTP, 10 HEPES, 10 EGTA, pH adjusted to 7.30 with CsOH to an osmolarity of 285 to 290 mOsm. To balance the osmolarity when EGTA was lowered to 0.5 mM, CsCH_3_SO_4_ was raised to 112 mM. The theoretical liquid junction potentials created between the extracellular recording solution and intracellular solutions were: 8.9 and 9.6 mV, for 10 and 0.5 mM EGTA, respectively. The voltage-clamp data presented here has not corrected for the junction potential; thus, the actual applied voltages are shift by ~ −9 to −10 mV from what is stated in the manuscript.

Whole-cell patch-clamp measurements were made with a HEKA EPC-10 amplifier equipped with Patchmaster software (Lambrecht, Pfalz, Germany). We used the software’s ‘sine+dc’ lock-in operation mode to monitor changes in membrane capacitance, conductance, and series resistance. Whole-cell electrodes were fabricated from thick-wall glass capillary tubes, and their tip region was coated with Sylgard (Dow Corning). Pipette resistance was 9 to 11 MOhms. The cell’s voltage was held at −70 mV around which a 2 kHz sine wave with a 50 mV peak-to-peak amplitude (−95 to −45 mV) was applied. The lock-in outputs were sampled at 20 kHz and filtered online with the low-pass f_c_ set to 2.9 kHz. The I-V_step_ protocols were sampled at 50 kHz and filtered online with the low-pass f_c_ set to 10 kHz.

Patch-clamp recordings in this study targeted rod soma in the OPL, which contain the ribbon in the soma compartment, referred to as the ‘soma-ribbon’ configuration (Hagiwara et al., 2018). Immediately before making the on-cell seal, the extracellular solution was exchanged to an MES with 2.0 mM Ca^2+^ and TEA/Cs/TBOA. After gaining whole-cell access, rods were held at a V_m_ of −70 mV. The cells were infused for 30 to 40 s before the evoked release protocols began, which entailed a sequence of 5 or 7 depolarizations with stimulations given at 8 s intervals. Evoked release was studied within the first 2 minutes, and then I-V protocols were performed afterwards. The passive electrical properties of the soma-ribbon measured in voltage-clamp were on average as follows, R_series_: 29.7 ± 0.6 MΩ and whole-cell capacitance C_m_: 1.02 ± 0.03 pF; yielding a membrane time constant τ_*RC*_ ~ 30 μsec (see Table S1 and Fig 1B). Recordings were made at an ambient temperature of 30 to 32°C. Almost all recordings were made within 30 to 45 minutes after slicing, and typically 1 to 2 successful recordings (cells) per mouse.

Data analysis. The evoked ΔC_m_ was assessed as outlined in Figure 1H and 1I. Segments of the C_m_ trace, 50 msec in length, before and after depolarization were averaged and the difference equaled the evoked ΔC_m_; ΔG_m_ and ΔG_s_ were calculated in the same way. Since G_m_ is influenced by Cl^−^-currents arising from Ca^2+^-activated TMEM16A/B channels and the glutamate transporter, the following precautions were taken. For experiments with 0.5 mM EGTA in the intracellular solution, the post-depolarization segment was averaged after I_Cl(Ca)_ relaxed, 75 msec after the end of the stimulation. Next, TFB-TBOA was used to block the glutamate transporter tail-currents (concentration described above). Finally, lock-in amplifier outputs were monitored between depolarization episodes, and this was done by taking 100 msec sine wave sweeps (Fig. 1H)., and the difference between 10 msec windows averaged at the start (time point: 5 to 15 msec) and end (t: 85 to 95 msec) of the 100 ms sweeps were treated as baseline/between stimulation Δ values (see Fig. 1E).

The I_Ca_ amplitude and activation time constant were determined by fitting the onset of the inward membrane current with a single exponential. When 0.5 or 10 mM EGTA was used, fits started after a 200 to 300 μsec delay from the start of V_step_, and the fit ended approximately 3 msec later (see Fig. 2Aiv). The exception was when 0.5 mM EGTA was used and V_step_’s were made more positive than −20 mV, which accelerated I_Cl(Ca)_ onset (Fig. 1Ci). This left 1 to 2 msec to fit peak-IC_a_ when V_step_ was −10 to +30 mV. The E_Cl_ is predicted to be −51 mV, and after adding the liquid junction potential (9.6 mV), the E_Cl_ was estimated to be ~ −41 mV. Finally, the I-V relationships were fitted with a Boltzmann equation: I=I_max_*(I_min_−I_max_)/(1+exp((V_m_−V_1/2_)/*k*)), and a modified form: I=G_max_*(V_m_−V_rev_)/(1+exp(−(V_m_-V_1/2_)/*k*)), where I is the peak current, V_m_ is the membrane voltage, V_1/2_ is the voltage for half activation, V_rev_ is the reversal potential, G_max_ is the maximal conductance, and *k* is the slope factor. Fitting and statistical analysis were performed with Origin software (OriginLab Corporation). All experimental values are given as mean ± SE.

### ERG recordings

Mice were dark adapted overnight. They were anesthetized with an intraperitoneal injection of ketamine (0.125 mg/g) and xylazine (2.5 μg/g), and pupils dilated with 1% atropine sulfate. A AgCl wire ring-electrode was placed on the cornea, and electrical contact was made with a NaCl saline solution, plus methylcellulose to maintain moisture. A needle reference electrode was inserted subcutaneously above the nose, and a ground electrode was inserted near the tail. A custom-designed Ganzfield illuminator with 25 white LEDs was used to deliver 0.1 msec light flashes every 5 sec, which were incrementally increased in intensity from 0.0003 to 0.278 cds/m^2^ (calibrated with a Mavolux, IPL 10530). Recorded potentials were amplified, low-pass filtered at 8 kHz, and sampled at a rate of 24 kHz. Ten responses were averaged per light intensity. The ergs were low-pass filtered using a FFT set to a corner-frequency of 400 Hz or 20 Hz for measurement of a- or b-wave parameters, respectively. All analysis were performed with Origin software (OriginLab Corporation).

### Converting ΔC_m_ into the number of fusing vesicles

To calculate the capacitance per vesicle we relied on published data. Single SV fusion events measured from mouse inner hair cells have an average C_m_: 40.2 aF and SV diameter of 36.9 nm (Grabner and Moser, 2018). Mouse rods have smaller SVs with an average diameter: 32.5 nm (Grabner et al., 2015). The calculated C_m_ per mouse rod SV is 35.4 aF.

## General

We would like to thank Stefan Thom for excellent technical assistance with the erg recordings. We thank Drs. Erwin Neher and Jakob Neef for critical feedback on the manuscript.

## Author contributions

Experiments were designed, performed and analyzed by C.P.G., and the manuscript was written by C.P.G. and T.M..

## Supplementary Materials

**Figure S1.**
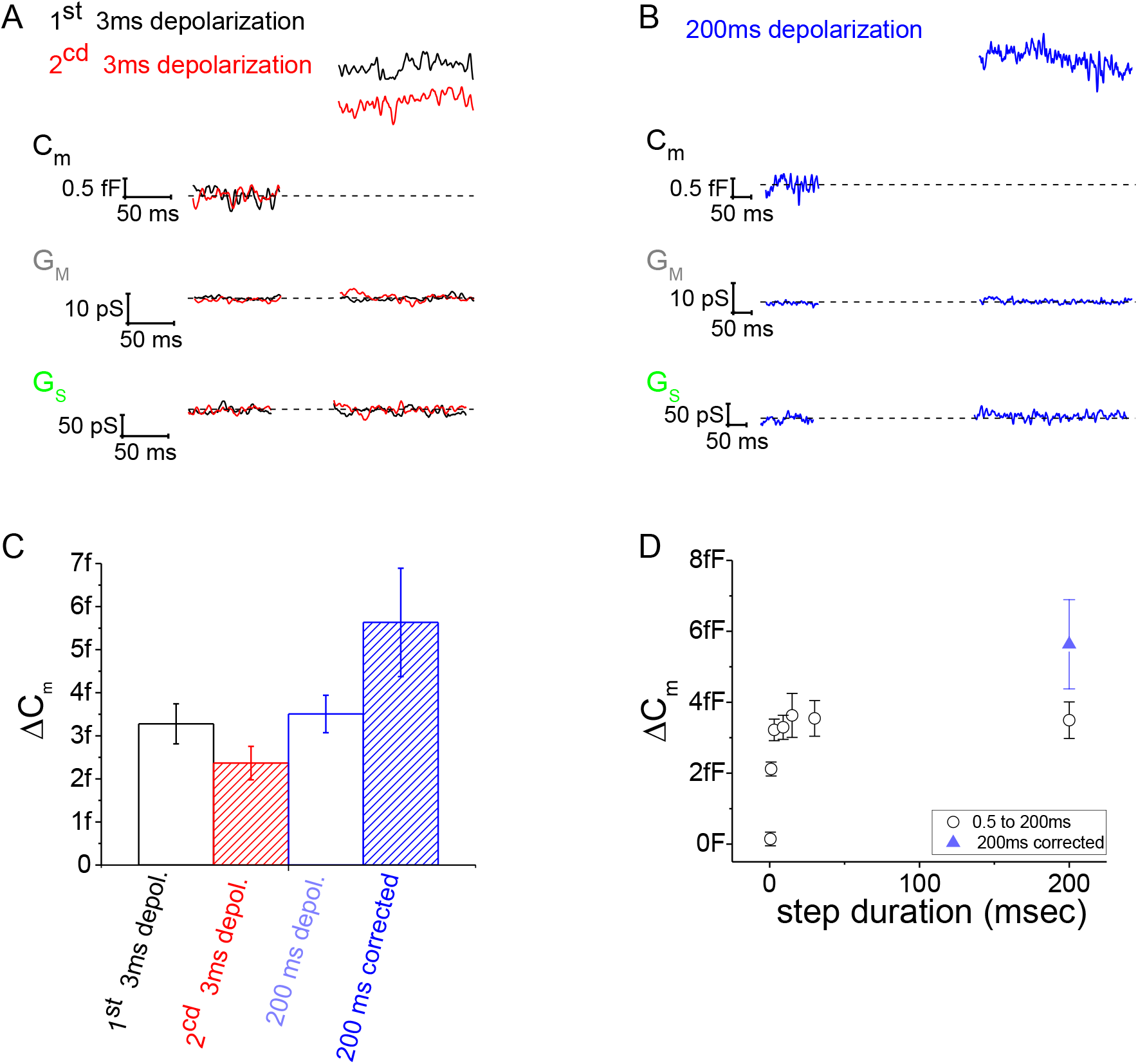
Estimation of the continual release rate. The degree of release run down was assessed by comparing 3 msec depolarizations (V_depol_ −18 mV) at different time points. The first 3 msec depolarization was given as part of a series of different step durations (as presented in Fig. 1E-J), at approximately 50 to 60 sec into the experiment. The second 3 msec depolarization was given at approximately 130 sec into the experiment, and it preceded the 200 msec depolarization by 12 sec. (**A**) The average lock-in output traces are presented for the 1^st^ and 2^cd^ 3 msec steps (7 cells). (**B**) The lock-in output traces are presented for the 200 msec steps (7 cells). (**C**) Presents the average ΔC_m_ results derived from 3 and 200 msec depolarizations. To calculate the 200 msec ΔC_m_ corrected for rundown within recording, the 200 msec ΔC_m_ was divided by the rundown factor: ((2^cd^ 3 msec ΔC_m_)/(1^st^ 3 msec ΔC_m_)).

**Figure S2.**
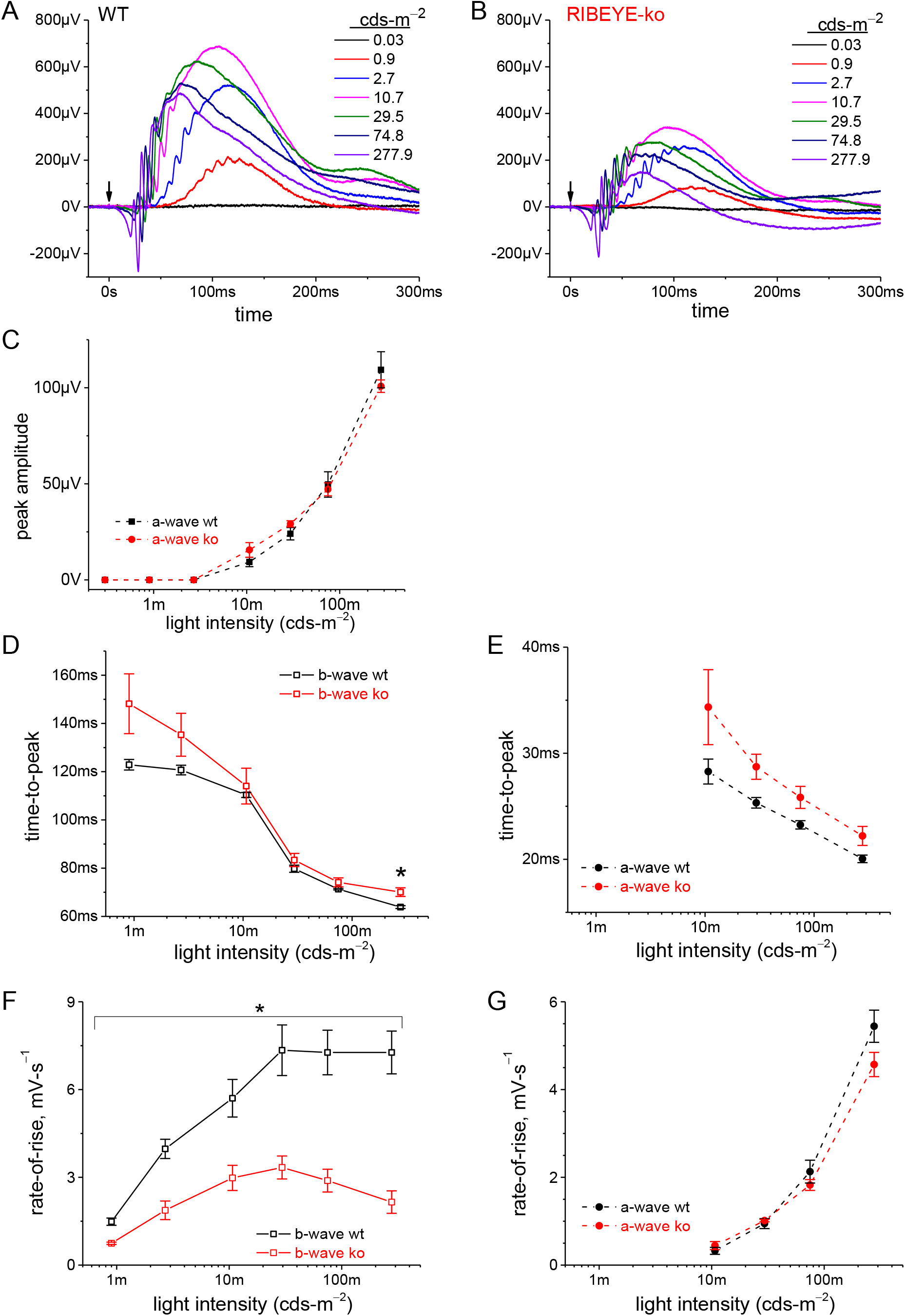
Rod driven light responses are suppressed in RIBEYE-ko mice. (**A**) Dark-adapted ergs recorded from an individual wt mouse. Intensity of the scotopic, dim light flashes are indicated in the graph. The arrow marks the moment of the 0.1 msec light flash. ERGs are presented at full band-width, without offline filtering. (**B**) Dark-adapted erg recording from an individual ko mouse. (**C**) Plot of erg a-wave amplitudes over the range of flash intensities, which showed no statistically significant difference between wt and ko mice. See Figure 4M for the b-wave responses. (**D**) As the flash intensity increased, the b-wave time-to-peak shortened. The only statistically significant difference between genotypes was witnessed at the highest flash intensity; *, p: 0.034. (**E**) As the flash intensity increased, the a-wave time-to-peak shortened. There were no statistically significant differences between genotypes. (**F**) WT b-waves rate-of-rise were significantly greater than ko mice across the full range of flash intensities; *, p-values: 0.04 to 0.009. (**G**) The a-wave rate-of-rise was not significantly different for the two genotypes. Average responses in C-G were attained from 4 ko and 3 wt mice.

**Table S1.** Whole-cell patch-clamp recording parameters.

|  | Lockin-Amplifier outputs |  |  |  |  |
| --- | --- | --- | --- | --- | --- |
| Ribeye Wt<br>0.5mM EGTA<br>(21 cells) | Cm<br>(Farads) | Gm<br>(Siemens) | Gs<br>(Siemens) | Ra<br>(MOhm) | Leak at<br>-70mV<br>(Ampere) |
| *Mean* | 974f | 67.7p | 28.3n | 31.3M | -4.83p |
| *SE of mean* | 39.8f | 11.5p | 2.03n | 3.74M | 928f |
| *Minimum* | 460f | 7.30p | 14.0n | 14.8M | -17.0p |
| *Median* | 987f | 45.0p | 28.0n | 21.5M | -3.20p |
| *Maximum* | 1.35p | 220p | 54.0n | 88.0M | 0.00 |
| Ribeye (-/-)<br>0.5mM EGTA<br>(17 cells) | Cm<br>(Farads) | Gm<br>(Siemens) | Gs<br>(Siemens) | Ra<br>(MOhm) | Leak at<br>-70mV<br>(Ampere) |
| *Mean* | 1.05p | 88.4p | 32.0n | 27.1M | -6.26p |
| *SE of mean* | 62.8f | 20.5p | 3.82n | 2.91M | 1.61p |
| *Minimum* | 679f | 15.1p | 11.2n | 11.7M | -20.0p |
| *Median* | 948f | 49.3p | 30.8n | 22.3M | -2.30p |
| *Maximum* | 1.67p | 306p | 71.7n | 57.7M | 0.00 |
| Ribeye Wt<br>10mM EGTA<br>(21 cells) | Cm<br>(Farads) | Gm<br>(Siemens) | Gs<br>(Siemens) | Ra<br>(MOhm) | Leak at<br>-70mV<br>(Ampere) |
| *Mean* | 1.01p | 58.3p | 23.2n | 29.0M | -3.35p |
| *SE of mean* | 20.4f | 8.65p | 1.23n | 3.64M | 839f |
| *Minimum* | 897f | 14.0p | 12.8n | 12.8M | -12.0p |
| *Median* | 995f | 54.1p | 23.5n | 20.9M | -3.00p |
| *Maximum* | 1.28p | 137p | 31.1n | 68.4M | 1.50p |
| Ribeye (-/-)<br>10mM EGTA<br>(10 cells) | Cm<br>(Farads) | Gm<br>(Siemens) | Gs<br>(Siemens) | Ra<br>(MOhm) | Leak at<br>-70mV<br>(Ampere) |
| *Mean* | 974f | 39.5p | 21.9n | 37.4M | -3.57p |
| *SE of mean* | 162f | 34.3p | 8.16n | 13.8M | 3.79p |
| *Minimum* | 665f | 10.4p | 9.99n | 18.9M | -12.0p |
| *Median* | 968f | 25.9p | 21.4n | 42.0M | -2.00p |
| *Maximum* | 1.21p | 111p | 37.1n | 51.7M | -1.00p |

**Table S2.**
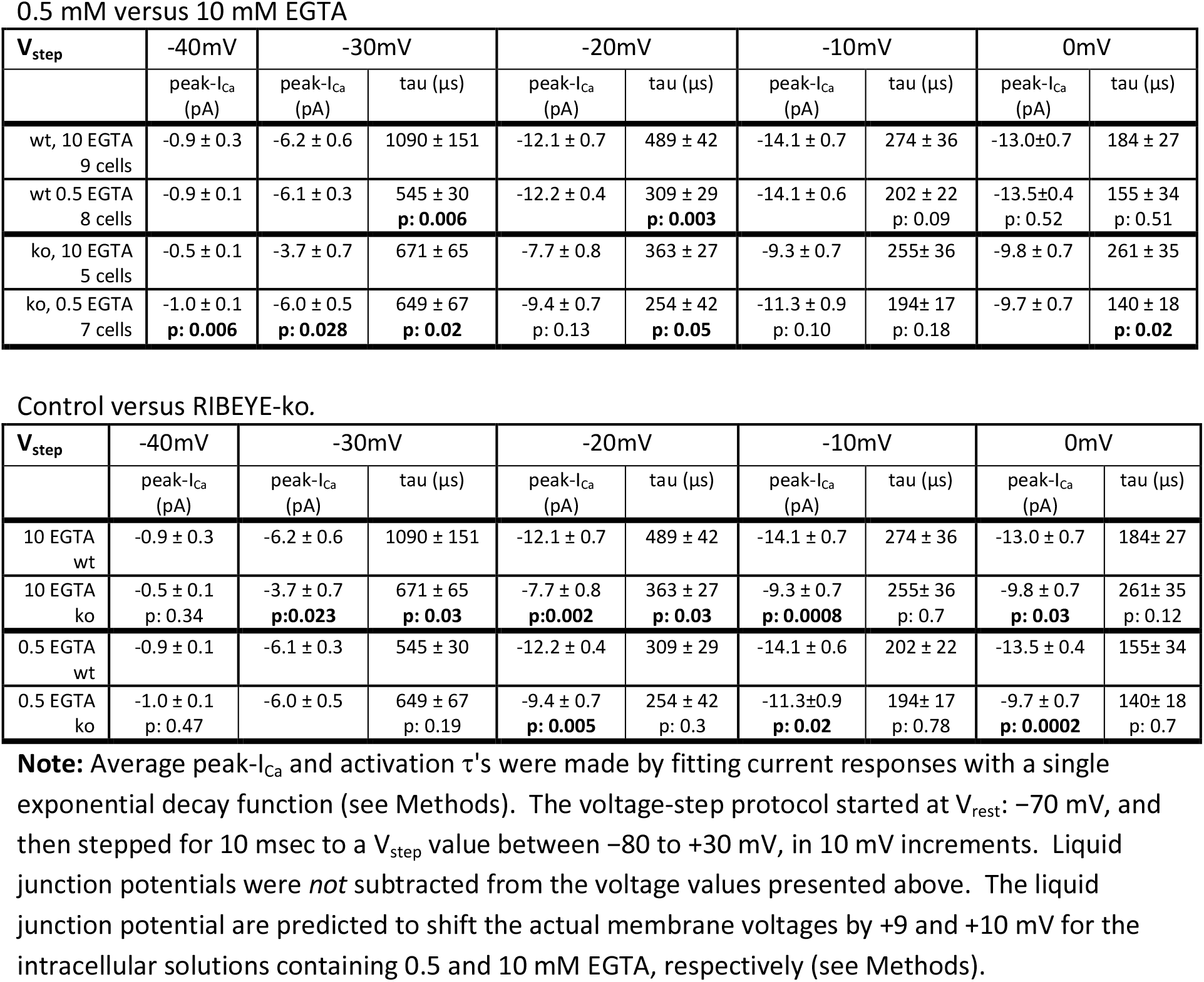
Activation kinetics for Ca_v_ currents.

**Table S3.**
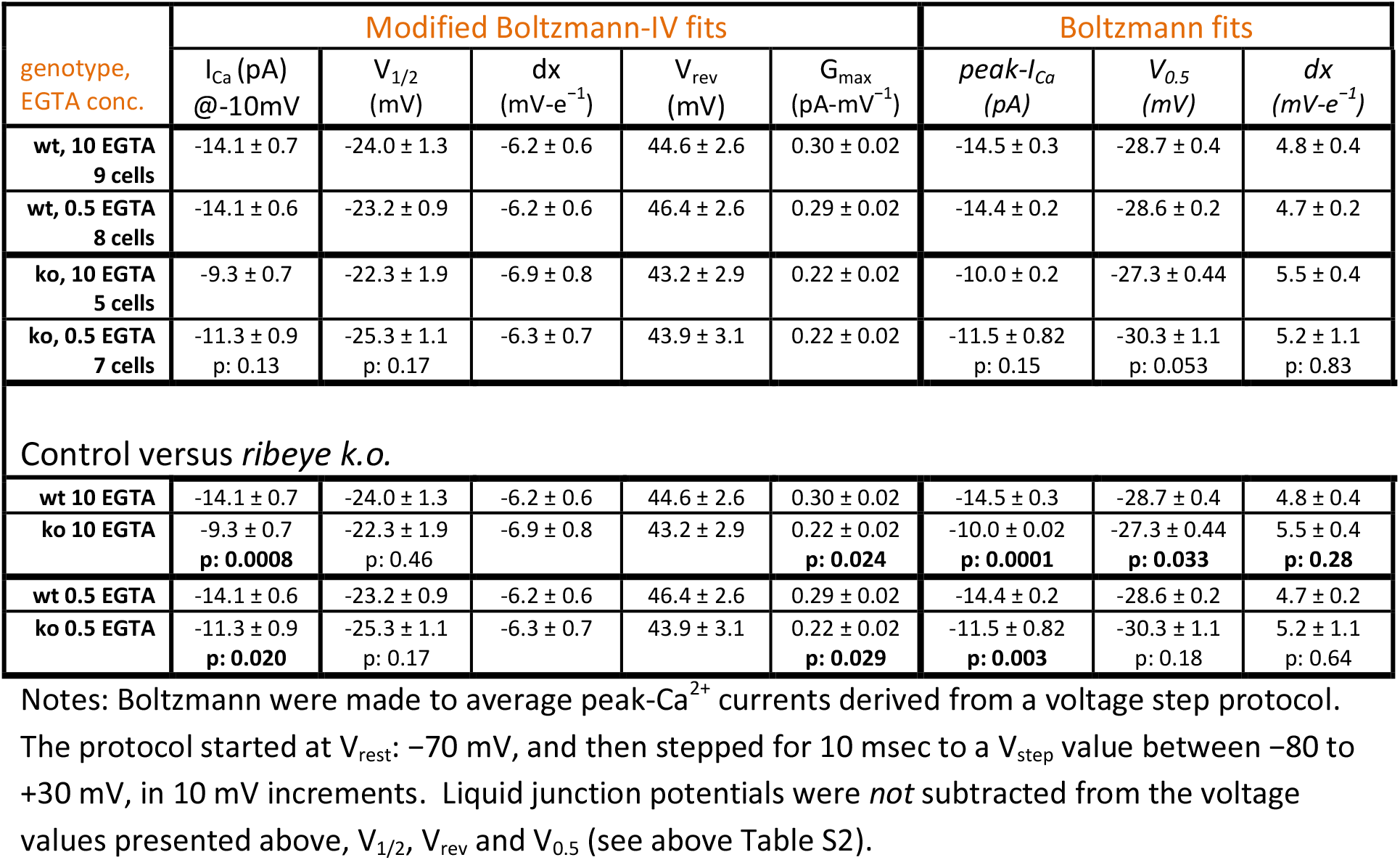
Peak-I_Ca_-Voltage relationship for wild type and RIBEYE-ko rods filled with 0.5 or 10 mM EGTA.

| genotype,<br>EGTA conc. | Modified Boltzmann-IV fits |  |  |  |  | Boltzmann fits |  |  |
| --- | --- | --- | --- | --- | --- | --- | --- | --- |
| | $I_{Ca}$ (pA)<br>@-10mV | $V_{1/2}$<br>(mV) | $dx$<br>(mV-$e^{-1}$) | $V_{rev}$<br>(mV) | $G_{max}$<br>(pA-mV$^{-1}$) | $peak-I_{Ca}$<br>(pA) | $V_{0.5}$<br>(mV) | $dx$<br>(mV-$e^{-1}$) |
| wt, 10 EGTA<br>9 cells | -14.1 ± 0.7 | -24.0 ± 1.3 | -6.2 ± 0.6 | 44.6 ± 2.6 | 0.30 ± 0.02 | -14.5 ± 0.3 | -28.7 ± 0.4 | 4.8 ± 0.4 |
| wt, 0.5 EGTA<br>8 cells | -14.1 ± 0.6 | -23.2 ± 0.9 | -6.2 ± 0.6 | 46.4 ± 2.6 | 0.29 ± 0.02 | -14.4 ± 0.2 | -28.6 ± 0.2 | 4.7 ± 0.2 |
| ko, 10 EGTA<br>5 cells | -9.3 ± 0.7 | -22.3 ± 1.9 | -6.9 ± 0.8 | 43.2 ± 2.9 | 0.22 ± 0.02 | -10.0 ± 0.2 | -27.3 ± 0.44 | 5.5 ± 0.4 |
| ko, 0.5 EGTA<br>7 cells | -11.3 ± 0.9<br>p: 0.13 | -25.3 ± 1.1<br>p: 0.17 | -6.3 ± 0.7 | 43.9 ± 3.1 | 0.22 ± 0.02 | -11.5 ± 0.82<br>p: 0.15 | -30.3 ± 1.1<br>p: 0.053 | 5.2 ± 1.1<br>p: 0.83 |
| Control versus *ribeye k.o.* |  |  |  |  |  |  |  |  |
| wt 10 EGTA | -14.1 ± 0.7 | -24.0 ± 1.3 | -6.2 ± 0.6 | 44.6 ± 2.6 | 0.30 ± 0.02 | -14.5 ± 0.3 | -28.7 ± 0.4 | 4.8 ± 0.4 |
| ko 10 EGTA | -9.3 ± 0.7<br>p: 0.0008 | -22.3 ± 1.9<br>p: 0.46 | -6.9 ± 0.8 | 43.2 ± 2.9 | 0.22 ± 0.02<br>p: 0.024 | -10.0 ± 0.02<br>p: 0.0001 | -27.3 ± 0.44<br>p: 0.033 | 5.5 ± 0.4<br>p: 0.28 |
| wt 0.5 EGTA | -14.1 ± 0.6 | -23.2 ± 0.9 | -6.2 ± 0.6 | 46.4 ± 2.6 | 0.29 ± 0.02 | -14.4 ± 0.2 | -28.6 ± 0.2 | 4.7 ± 0.2 |
| ko 0.5 EGTA | -11.3 ± 0.9<br>p: 0.020 | -25.3 ± 1.1<br>p: 0.17 | -6.3 ± 0.7 | 43.9 ± 3.1 | 0.22 ± 0.02<br>p: 0.029 | -11.5 ± 0.82<br>p: 0.003 | -30.3 ± 1.1<br>p: 0.18 | 5.2 ± 1.1<br>p: 0.64 |

Notes: Boltzmann were made to average peak- $Ca^{2+}$ currents derived from a voltage step protocol.

**Table S4.** Ca^2+^-dependent tail-currents measured with 0.5 mM EGTA in the pipette.

| genotype | I<sub>Cl(Ca)</sub> @<br>-10 mV<br>(pA) | Modified Boltzmann-IV fits |  |  |  | Boltzmann fits |  |  |
| --- | --- | --- | --- | --- | --- | --- | --- | --- |
|  |  | V<sub>1/2</sub><br>(mV) | dx<br>(mV-e<sup>-1</sup>) | V<sub>rev</sub><br>(mV) | G<sub>max</sub><br>(pA-mV<sup>-1</sup>) | span<br>(pA) | V<sub>0.5</sub><br>(mV) | dx<br>(mV-e<sup>-1</sup>) |
| wt 0.5 EGTA<br>n: 6 | -68 ± 13 | -20.4 ± 0.5 | -5.55 ± 0.95 | 35.8 ± 3.8 | 1.41 ± 0.44 | 67 ± 14 | -23.9 ± 0.8 | 3.20 ± 0.19 |
| ko 0.5 EGTA<br>n: 5 | -27 ± 5<br>p: 0.036 | -22.8 ± 0.6<br>p: 0.016 | -5.28 ± 0.25<br>p: 0.79 | 34.1 ± 3.0<br>p: 0.73 | 0.70 ± 0.13<br>p: 0.17 | 27 ± 5<br>p: 0.042 | -27.4 ± 0.8<br>p: 0.013 | 3.52 ± 0.24<br>p: 0.33 |

Notes: Ca<sup>2+</sup>-activated tail-currents were measured following the voltage step protocol used to measure peak-I<sub>Ca</sub> amplitude. The protocol started at V<sub>rest</sub>: -70 mV, and then stepped for 10 msec to a V<sub>step</sub> value between -80 to +30 mV, in 10 mV increments. Tail-currents were measured 3 msec after repolarizing the cell to -70 mV. Liquid junction potentials were *not* subtracted from the voltage values presented above, V<sub>1/2</sub>, V<sub>rev</sub> and V<sub>0.5</sub> (see Table 2 for LJP).

